# Colour polymorphism associated with a gene duplication in male wood tiger moths

**DOI:** 10.1101/2022.04.29.490025

**Authors:** Melanie N. Brien, Anna Orteu, Eugenie C. Yen, Juan A. Galarza, Jimi Kirvesoja, Hannu Pakkanen, Kazumasa Wakamatsu, Chris D. Jiggins, Johanna Mappes

## Abstract

Colour is often used as an aposematic warning signal, with predator learning expected to lead to a single colour pattern within a population. However, there are many puzzling cases where aposematic signals are also polymorphic. The wood tiger moth, *Arctia plantaginis*, displays bright hindwing colours associated with unpalatability, and males have discrete colour morphs which vary in frequency between localities. In Finland, both white and yellow morphs can be found, and these colour morphs also differ in behavioural and life-history traits. Here, we show that male colour is linked to an extra copy of a *yellow* family gene that is only present in the white morphs. This white-specific duplication, which we name *valkea*, is highly upregulated during wing development, and CRISPR knockouts validate the role of *valkea* in producing white wing colour. We also characterise the pigments responsible for yellow, white and black colouration, showing that yellow is partly produced by pheomelanins, while black is dopamine-derived eumelanin. Our results add to a growing number of studies on the genetic architecture of complex and seemingly paradoxical polymorphisms, and the role of gene duplications and structural variation in adaptive evolution.

## Introduction

Colour polymorphisms, defined as the presence of multiple discrete colour phenotypes within a population (Huxley, 1955), provide an ideal trait to study natural and sexual selection. Colour phenotypes can have an effect on fitness in many contexts including camouflage, mimicry and mating success. Colour is often associated with aposematism, where it acts as a signal, warning predators of unpalatability (Cott, 1940; Cuthill *et al*., 2017). In such cases, predator learning should favour the most common colour pattern, leading to positive frequency dependent selection (Endler, 1988). Despite this, aposematic polymorphisms can be stable when selection is context-dependent (Briolat *et al*., 2019), especially where genetic correlations between colour phenotypes and other traits lead to complex fitness landscapes (reviewed by McKinnon and Pierotti, 2010).

A variety of genetic mechanisms can underpin these types of complex polymorphisms involving multiple associated traits (Orteu and Jiggins, 2020). In many cases, such complex polymorphisms are controlled by ‘supergenes’ in which divergent alleles at several linked genes are maintained in strong linkage disequilibrium by reduced recombination. The most common mechanism for locally reduced recombination are inversions, which range from single inversions involving a small number of genes, to multiple nested inversions covering large genomic regions (Joron *et al*., 2011; Wang *et al*., 2013; Küpper *et al*., 2016; Funk *et al*., 2021). Nonetheless, other mechanisms for reducing recombination, such as centromeres or large genomic deletions, may also play a role. An alternative mechanism is that a single regulatory gene controls variation via multiple downstream effects (Thompson and Jiggins, 2014). While there are fewer instances in which multiple phenotypes seem to be controlled by a single gene, one potential example is the common wall lizard, where colour genes have pleiotropic effects on behavioural and reproductive traits (Andrade *et al*., 2019). Multiple mutations within a single gene can also lead to variation in multiple traits (Linnen *et al*., 2013).

The wood tiger moth, *Arctia plantaginis*, has a complex polymorphism that has been well studied in an ecological context. Males show polymorphic aposematic hindwing colouration with discrete yellow, white or red hindwing colour morphs found at varying frequencies in different geographic locations. In Finland, for example, both yellow and white morphs can be found, with white morphs varying in frequency from 40 to 75% (Galarza *et al*., 2014). In Estonia, white morphs make up 97% of the population, while yellows morphs form a completely monomorphic population in Scotland (Hegna, Galarza and Mappes, 2015) (Figure 1). Long term breeding studies of these moths have shown that male hindwing colour is a Mendelian trait controlled by a single locus with two alleles (Suomalainen, 1938; Nokelainen *et al*., 2022). White alleles (W) are dominant over the yellow (y). These colour genotypes also covary with behavioural and life-history traits, contributing to the maintenance of this polymorphism. Yellow males are subject to lower levels of predation in the wild (Nokelainen *et al*., 2012, 2014), while white males have a positive frequency-dependent mating advantage (Gordon *et al*., 2015). There are differences in chemical defences, and bird reactions to these defences, between the genotypes (Rojas *et al*., 2017; Winters *et al*., 2021). Yellow morphs show reduced flight activity compared to white males, although yellows may fly at more selective times, i.e. at peak female calling periods (Rojas, Gordon and Mappes, 2015). Increased reproductive success in Wy genotype females points towards strong heterozygote advantage (De Pasqual *et al*., 2022). In summary, there is a trade-off between natural selection through predation and reproductive success, which contributes to the maintenance of this polymorphism (Rönkä *et al*., 2020).

**Figure 1:**
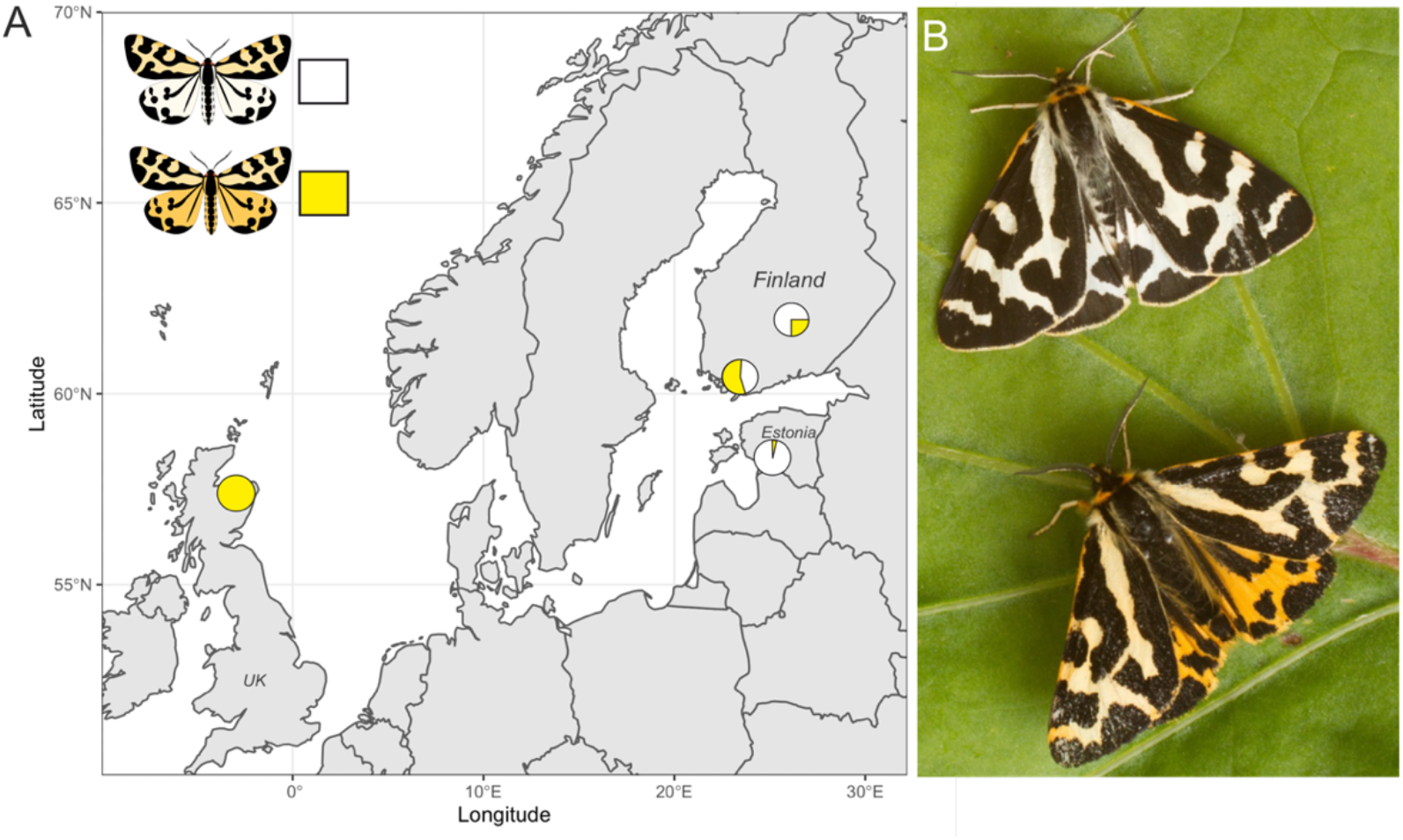
(A) Sampling locations and frequencies of yellow and white *Arctia plantaginis* males in Finland, Scotland and Estonia. (B) Males of the white and yellow colour morphs (Credit: Samuel Waldron).

Despite the large body of research on *A. plantaginis* colour morphs, the genetic basis of this polymorphism is unknown. Here, we explore male hindwing colour variation using linkage mapping, whole genome data and gene expression analyses, with wild populations and lab crosses, to identify the locus controlling the colour polymorphism. We use CRISPR/Cas9 gene knockouts to determine the function of the identified gene, and then characterise the pigments producing yellow, white, and black colouration on the wings of male *A. plantaginis*. Our findings aim to provide an example of the genetic architecture controlling a trait that is part of a complex polymorphism.

## Results

### A narrow genomic region is associated with hindwing colour

To investigate the genetic basis of male hindwing colouration in *A. plantaginis*, we carried out a quantitative trait locus (QTL) mapping analysis using crosses between heterozygous Wy males and homozygous yy females. We used RADseq data aligned to the yellow *A. plantaginis* reference genome from 172 male offspring (90 white and 82 yellow) from four families. The QTL analysis identified a single marker associated with male hindwing colour (Figure 2A). This marker was found on scaffold YY_tarseq_206_arrow at position 9,887,968bp (95% confidence intervals 9,349,978-9,888,009bp) and had a LOD score of 32.8 (p<0.001). The significant marker explains around 75% of the phenotypic variation and, with one exception, yellow individuals all had a homozygous yy genotype at this marker.

**Figure 2:**
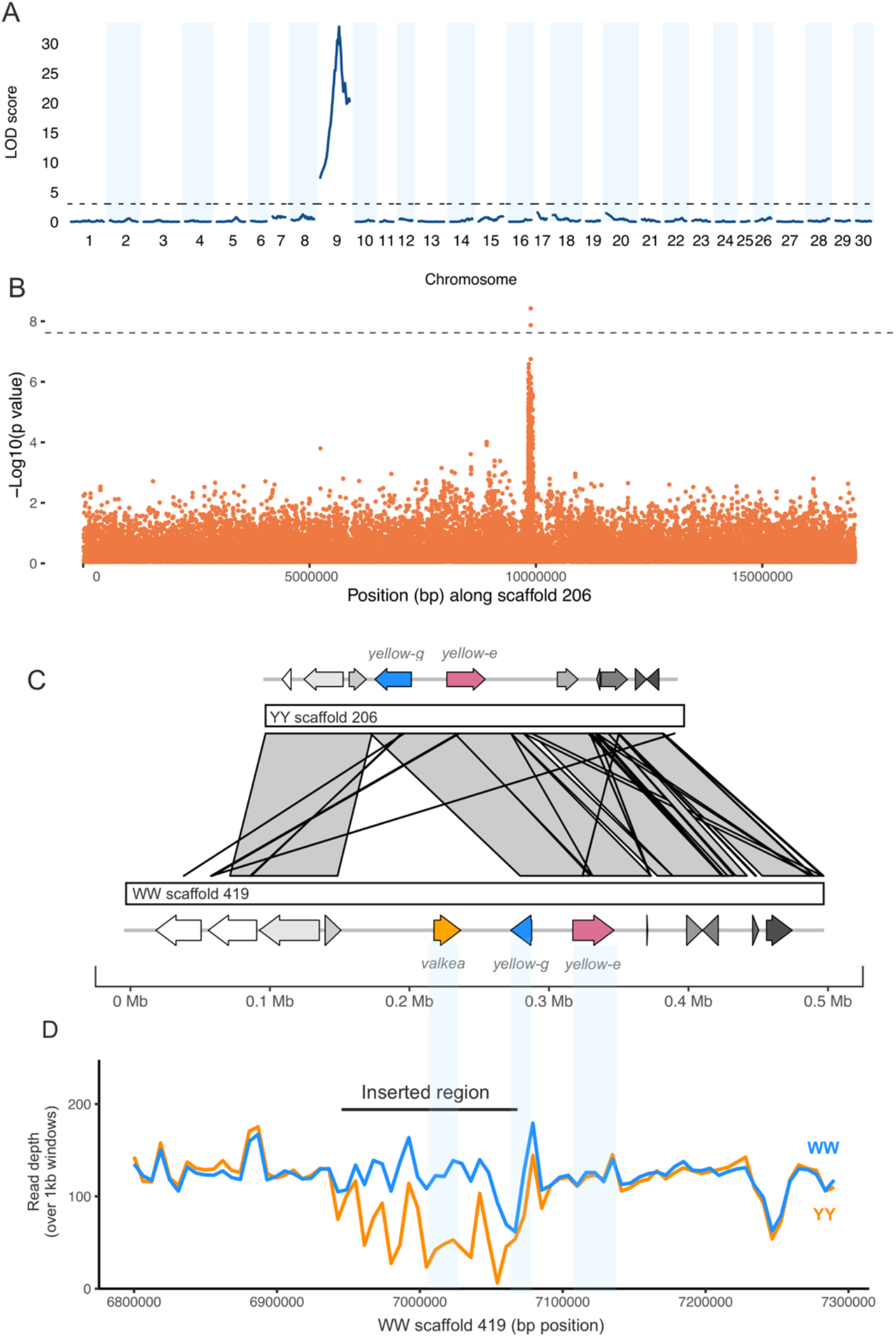
A duplicated region in white morphs is associated with male hindwing colour. (A) QTL analysis of white and yellow F1 males (n=172) reveals a 500kb region significant on scaffold 206, part of linkage group 9. The dotted line indicates the significance threshold determined by permutation tests (p=0.05). (B) A GWAS of wild samples (n=46) showed SNPs associated with hindwing colour along the same scaffold. The dotted line shows the Bonferroni corrected significance threshold. (C) An insertion in the white reference sequence contains a copy of the *yellow-e* gene, which we named *valkea*, in addition to the *yellow-e* present in both white and yellow morphs. (D) Mean read depth across the candidate region in all Finnish white (WW and Wy) and yellow (yy) samples.

To further narrow down this region, we ran a genome wide association study (GWAS) using whole genome sequences of males from four populations: polymorphic Southern Finland (5 white, 5 yellow) and Central Finland (10 white, 10 yellow), Estonia (4 white), where males are mostly white, and monomorphic Scotland (12 yellow), where all males are yellow. This identified a region of associated SNPs also on scaffold 206 (Figure 2B). Two SNPs, 137bp apart (at positions 9,885,384 and 9,885,521), were significant above a strict Bonferroni corrected threshold. 162 SNPs were over the threshold of p<0.0001 and, of these, 155 are within a 99kb region on scaffold 206 (9,833,387-9,932,264bp). The top SNPs are within 2.5kb from the top QTL marker, and the SNP at this marker has a p-value <0.0001.

The 538Kb QTL interval contains 21 genes (Table S1) which were annotated with reference to *D. melanogaster*. Of these genes, four are part of the *yellow* gene family. The top two SNPs from the GWAS, and the top marker from the QTL, fall in a non-coding region upstream of the gene, *yellow-e*, and are also close to an additional *yellow* gene, *yellow-g*.

### Identifying structural variation in this region

The trio binning method used by Yen et al. (2020) to assemble the *A. plantaginis* reference genome produced two reference sequences, one for a white allele and one for a yellow allele. We extracted the region containing the QTL interval from the yellow reference and aligned it against the white reference. The alignment showed a duplicated region approximately 117kb long on scaffold 419 of the white reference from around 6,941,000-7,058,000bp (Figure 2 - figure supplement 1). The *yellow-e* gene and its flanking regions are within this sequence and are therefore duplicated in the white reference (Figure 2C). One copy of the gene (named jg1310 in the W annotation) has 7 exons and is similar to the *yellow-e* gene in the yellow reference (99.7% identity in coding sequences). The second copy unique to the white scaffold (jg1308) has only the first 5 exons (81.8% identical to the gene in the yellow reference), possibly due to a stop codon mutation in the 5^th^ exon. For clarity, we named this duplicated white-specific copy *valkea*, in reference to a Finnish word for ‘white’. While all white samples had consistent coverage of reads across the duplicated region, coverage was patchy in yellow samples, with many regions having no and very low coverage in yellow samples (Figure 2D). Those reads that map in the *valkea* region in yellow samples are likely to be mapping errors, because the sequence similarity is high and mapping quality is reduced within the duplication (Figure 2 – figure supplement 2). When increasing the mapping quality filtering, read depth decreases more in yellow samples compared to white samples in this region (Figure 2 – figure supplement 3). We confirmed the absence of this region in yellow individuals by designing primers within the duplication (Table S2, Appendix Figure 1), which only amplified in WW and Wy samples, including Finnish, Estonian and lab populations.

To confirm that both of these gene copies are related to *yellow-e*, we compared them to *yellow-e* orthologs found in *Bombyx mori, Heliconius melpomene* and *Drosophila melanogaster*, along with other *yellow* genes from *A. plantaginis* and *B. mori*. Both of the tiger moth genes were most closely related to the *H. melpomene yellow-e* (Figure 2 – figure supplement 4). Between *valkea* and *yellow-e* there is an additional gene which showed highest similarity to *Drosophila yellow-g2* (when extracted from both the white and yellow references). This gene is not part of the duplicated sequence and is present as a single copy in both morphs. Coverage across *yellow-g* and *yellow-e* genome regions in wild samples is similar in both morphs (Figure 2). Upstream of the duplication is an unnamed gene (listed as jg6744 in the yellow annotation and jg1307 in the white). This is the same orthologous gene in both reference genomes, having 99.3% identity. Similarly, if we look at the 150kb upstream region, sequence identity is 99.98%. There are no non-synonymous mutations between coding sequences of *yellow-e* when comparing the white and yellow references, although there are differences in the first exon of *yellow-g*. The absence of this duplicated region in the yellow morphs means we cannot determine if there is a change in linkage disequilibrium across white and yellow morphs.

### Valkea is differentially expressed between morphs

To pinpoint which of these candidate genes is associated with male wing polymorphism in *A. plantaginis*, we next performed gene expression analyses across several developmental stages. Based on knowledge of the expression patterns of *yellow* genes (Ferguson *et al*., 2011) and other melanin pathway genes such as *pale, ebony and ddc* in Lepidoptera (Zhang *et al*., 2017), we hypothesised that changes in gene regulation that control the development of wing colour morphs in the tiger moth most likely occur during pupal development. Pupal development in the wood tiger moth lasts for approximately 8 days, and no colour is present in the wings until day 7, when the yellow pigment appears. A few hours later, black melanin pigmentation is deposited. We sampled two stages early in development when no colouration is present in the wings (72 hours post-pupation, and 5 day old pupae), and two stages later in development: the point when yellow appears in yy morphs (Pre-mel, 7 day old pupae) and the other after black melanin has also been deposited (Mel, 7-8 days old). Forty individuals were sampled in total - five per genotype and stage.

First, we explored the general patterns of expression by mapping RNAseq reads to the white reference genome, which contains the duplicated region that includes *valkea*. We filtered out lowly expressed genes, retaining 10,920 genes and used Multidimensional Scaling (MDS), a dimensionality reduction technique, to explore which factors explain genome-wide variation in gene expression between samples. We observed that samples clustered based on their developmental stage, suggesting it is an important factor driving differences in genome-wide gene expression between samples (Figure 3 – figure supplement 1). Such a pattern would be expected as many genes are involved in development and thus are likely to be differentially expressed (DE) between developmental stages. No apparent clustering can be observed among samples of the same colour morph.

We next compared gene expression between yy and WW individuals at each of the developmental stages. Overall, 99 genes were differentially expressed (FDR < 0.05) between the two morphs (Figure 3A). Two of these DE genes, *yellow-e* and *valkea*, are two of the 22 genes identified in the GWAS and QTL analysis. *Valkea* was overexpressed in white individuals in the pre-melanin stage with a Log Fold Change of 10.32 and a p-value of 2.18e-06. As *valkea* is only fully present in the W genome, it is not expected to be expressed at all in the Y genome. *Yellow-e* was also overexpressed in white individuals during the pre-melanin stage with a Log Fold Change of 3.86 and adjusted p-value of 5.62e-06. In other developmental stages, neither *valkea* nor *yellow-e* showed differences in expression between morphs (Figure 3B).

**Figure 3:**
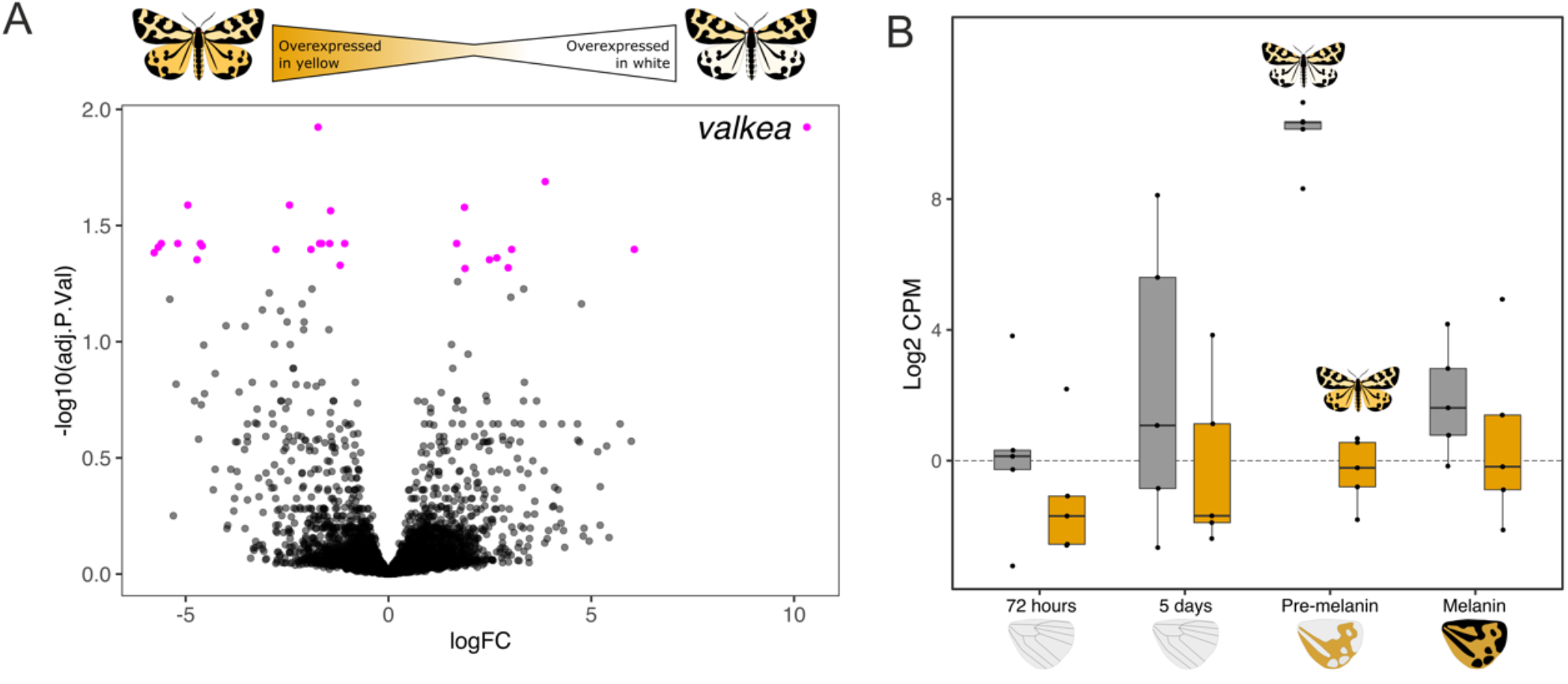
*Valkea* is overexpressed in white males in the pre-melanin stage. (A) In pink are genes that are significantly differentially expressed between yellow and white morphs at the pre-melanin stage. *Valkea* is the most DE gene (i.e. gene with the highest Log Fold Change). (B) Expression of *valkea* across developmental timepoints shows that it has higher expression measured in Log2 CPM (Counts Per Million) in white individuals compared to yellow ones. Expression of *valkea* in yellow morphs is around 0.

Of the 99 genes differentially expressed between white and yellow individuals across development, 49 were upregulated in the yy morph, while the remaining 50 were upregulated in WW individuals. The earliest developmental stage, 72-hours, was the stage with the highest number of DE genes (n=48), while the 5-day old stage had the fewest (n=7). One gene which encodes a C2H2 zinc finger transcription factor in *D. melanogaster*, ‘*jg15945*’, was over-expressed in yy in the first three stages (Figure 3 – figure supplement 2).

Finally, the GWAS and QTL peaks of association are situated in scaffold 419 of the WW reference assembly, which in a WW linkage map forms a linkage group along with 6 more scaffolds (472, 487, 515, 531, 540 and 609). We found that 12 genes present in this linkage group were differentially expressed, including *valkea* and *yellow-e* (Table S3), and identified their orthologues in *Drosophila melanogaster*.

### CRISPR/Cas9 knockouts of valkea produce yellow hindwings

To confirm the function of *valkea* in wing colouration, we used CRISPR/Cas9 to knock out the gene in white morphs. We tested 5 different guides to target the first 3 exons of *valkea* and injected Cas9/sgRNA duplexes into a total of 1223 eggs. Of 143 larvae that hatched only 6 developed to adults (Table S4). However, of the five males that did eclose, four had a visible change in phenotype. Males produced yellow scales instead of white on the dorsal side of both the forewings and the hindwings (Figure 4). Forewings were more yellow than in the wildtype yellow males, which usually have lighter forewings compared to hindwings. White scales on the ventral side of the wings also became yellow, similar to wildtype yellows. Black melanin patterning did not seem to be affected. Variation in the amount of melanin can be attributed to the populations from which the individuals originated, with the darker samples coming from the Finnish population (Figure 4 – figure supplement 1). Wildtype white morphs also reflect UV, particularly on the hindwings, but this is not seen in the CRISPR males (Figure 4 – figure supplement 2). This could suggest a change in scale structure, or that the yellow pigment covers the UV-reflecting structures.

**Figure 4:**
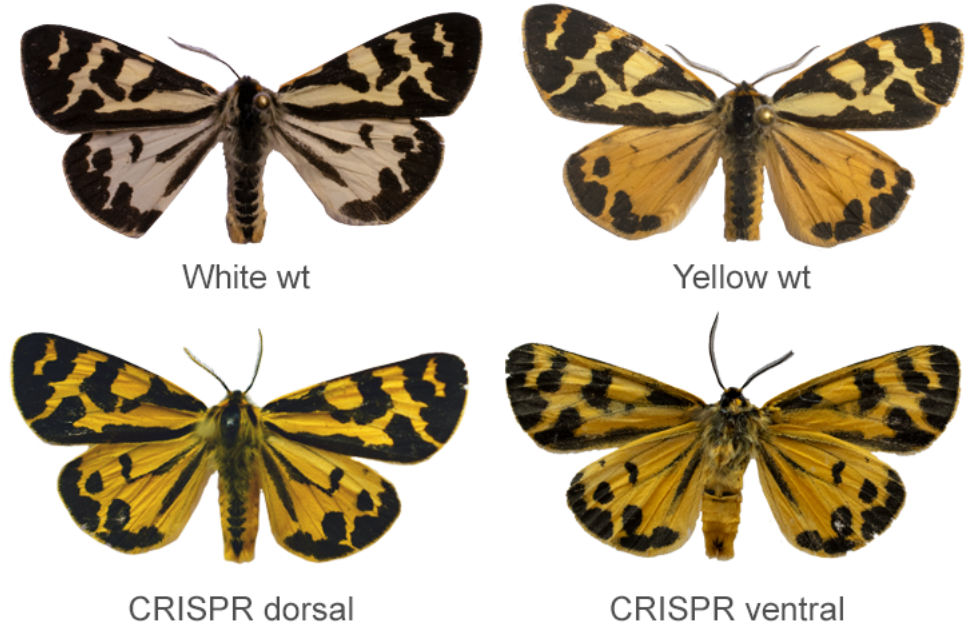
CRISPR/Cas9 knockouts of *valkea* transforms white scales into yellow scales across both hindwings and forewings. Wildtype WW and yy morphs (top), and the dorsal and ventral sides of one of the CRISPR knockout males (bottom).

Four out of the five guides tested produced a mutant phenotype, with no differences in the male phenotype between guides. The similarity of *valkea* to *yellow-e* means that it is likely that *yellow-e* could also have been knocked-out in these individuals. We used whole genome sequences of the mutants to confirm that the correct sites in *valkea* had been targeted (Figure 4 – figure supplement 3). All samples also showed evidence of editing at the corresponding *yellow-e* exons, which mainly involved insertions. The change in forewing colour could be attributed to a *yellow-e* mutation. Only one female survived to adulthood, and this had a mosaic phenotype. This individual had one yellow forewing, similar to the male mutants, with the rest of the body and wings being wildtype (Figure 4 – figure supplement 1). Since we did not expect a *valkea* knockout to affect female phenotypes as they always have red hindwings, this could be further evidence for the effect of *yellow-e* on forewing colour.

Survival of the eggs varied between the guides, although this was largely affected by the female, as hatching rate between females ranged from 0 to 70%. Females often lay unfertilised eggs so we expect that hatching rate will be low in some crosses. Using two guides in combination did not produce any pupae or adults.

### Pigment analysis

Since the *yellow* gene family, to which *valkea* is related, is known to be responsible for the production of melanin pigments, we further investigated the identity of the wing pigmentation. First, we ruled out the presence of several non-melanin pigment types in the hindwings, including pterins and carotenoids. Pterins are commonly found in insects and, along with purine derivatives, papiliochromes and flavonoids, are soluble in strong acids and bases or in organic solvents (Umebachi, 1975; Kayser, 1985; Shamim *et al*., 2014). We placed wing samples from each morph in NaOH overnight, then measured the absorbance of the supernatant using a spectrophotometer. We also left wings in methanol overnight before measuring the supernatant. The spectra did not show any peaks indicative of any pigment dissolved in the sample. Similarly, we found no evidence for carotenoid pigments after dissolving in a hexane:tert-butyl methyl ether solution (Appendix Figure 2). Wings did not fluoresce under UV light, providing further evidence for the lack of fluorescent pigments including pterins, flavonoids, flavins and papiliochromes (Umebachi, 1975; Kayser, 1985). Ommochromes are red and yellow pigments; high performance liquid chromatography (HPLC) ruled out the presence of these pigments on the moth wings, which we compared to data from ommochrome-containing *Heliconius* wings and a Xanthurenic acid standard (Appendix Figure 3).

HPLC analysis showed peaks characteristic of pheomelanin (Appendix Figure 4). Pheomelanins produce red-brown colour in grasshoppers and wasps (Galván *et al*., 2015; Jorge García, Polidori and Nieves-Aldrey, 2016), and orange-red colours in ants and bumblebees (Hines *et al*., 2017; Polidori, Jorge and Ornosa, 2017). Insects generally have dopamine-derived pheomelanin and a breakdown product of this is 4-Amino-3-hydroxyphenylethylamine (4-AHPEA) (Barek *et al*., 2018). Yellow wings showed large peaks for 4-AHPEA. White wings had around 27% of the 4-AHPEA levels seen in yellow wings, and black sections of the wings had 16%. Hydrogen iodide hydrolysis of wings produced the isomer 3-AHPEA, which may come from 3-nitrotyramine originating from the decarboxylation of 3-nitrotyrosine. Reduction of 3-nitrotyrosine produces 3-AHP, another marker of pheomelanin (Wakamatsu, Ito and Rees, 2002).

Analysis of the black portions of the wing found pyrrole-2,3-dicarboxylic acid (PDCA) and pyrrole-2,3,5-tricarboxylic acid (PTCA) (Appendix Figure 5). Both are components of eumelanin (Barek *et al*., 2018), suggesting that the black colouration seen in the wood tiger moth is predominantly eumelanin derived from dopamine. This is common in producing black colouration and providing structural components of the exoskeleton. In addition, dopamine is acylated to both *N*-β-alanyldopamine (NBAD) and *N*-acetyldopamine (NADA) sclerotins. NADA sclerotins are colourless and likely to be present on the white wings. This analysis of pigmentation is therefore consistent with a role for *yellow* family genes in regulating the colour polymorphism.

## Discussion

Hindwing colouration of male *Arctia plantaginis* is polymorphic and these colour morphs vary in multiple behavioural and life-history traits, providing an example of a complex polymorphism. Here, we have shown that variation in male hindwing colour is associated with a duplicated sequence found only in white morphs and containing a gene from the *yellow* gene family. The white-specific copy, *valkea*, is highly expressed during pupal development, consistent with genetic dominance of the white allele. When *valkea* is knocked out in the white morphs, yellow pigment is produced, providing functional evidence that *valkea* is responsible for the white phenotype rather than other genes in this region.

These results add to the increasing evidence for the role of gene duplications in the evolution of adaptive genetic variation. Genes for the metabolisms of proteins in *Heliconius* butterflies underwent several duplications, facilitating changes in diet and adaptation to pollen feeding (Smith, Macias-Muñoz and Briscoe, 2016). In *Zerene cesoina* butterflies, recent partial duplications of the transcription factor *doublesex*, resulting from multiple duplication events, are associated with sex-specific wing patterning. The duplicated paralog acts as a repressor of genes producing UV-reflecting wing scales in females (Rodriguez-Caro *et al*., 2021).

We hypothesise that the morph-specific duplication that we see in *A. plantaginis* provides a region of reduced recombination between morphs, as the duplicated region is effectively hemizygous and cannot recombine except in homozygote genotypes, which could contribute to the maintenance of the complex polymorphism and the linkage of multiple traits. This is similar to the genetic architecture of the *Primula* supergene controlling heterostyly, which involves a large duplication containing five genes (Huu *et al*., 2020). In polymorphic *Papilio dardanus*, one colour pattern morph is associated with a duplicated region, again providing physical constraints on recombination (Timmermans *et al*., 2014). Nonetheless, in the case of the wood tiger moth, it remains unclear how a single gene, such as *valkea*, can control the development of a broad array of phenotypic traits.

One possible mechanism is that there is a regulatory element along the scaffold which is controlling colour via the *valkea* gene, but also regulating other genes to control different phenotypic traits. We found the most significant markers and SNPs located in a non-coding region close to the *yellow* genes, which likely contains a cis-regulatory element (CRE) controlling transcription of *valkea*. In cichlids, for example, a CRE at the gene encoding agouti-related peptide 2 controls variation in strip patterning in two closely related species (Kratochwil *et al*., 2018). Conserved CREs were shown to have wide ranging effects on wing patterning in multiple Nymphalidae butterflies (Mazo-Vargas *et al*., 2022).

Differential expression of other genes on the same chromosome controlled by the CRE could explain variation in covarying traits. The over-expression of another gene, possibly encoding a zinc transcription factor, in yellow individuals in the early pupal stages suggests that there is differential expression of unlinked genes as a result of the polymorphism, although since this gene is on a different chromosome to *valkea* it is unlikely to be directly controlled by the CRE. Another hypothesis is that somehow *valkea* itself regulates other genes. However, *yellow* family genes are not known to regulate transcription of other genes, unlike for example *doublesex*, which undergoes alternative splicing and female mimetic wing pattern polymorphism in *Papilio polytes* (Kunte *et al*., 2014; Nishikawa *et al*., 2015).

The *yellow* family genes are highly conserved throughout insects (Ferguson *et al*., 2011). They have been widely linked to colouration (Wittkopp, Vaccaro and Carroll, 2002; Miyazaki *et al*., 2014; Zhang *et al*., 2017; Zhang, Mazo-Vargas and Reed, 2017), as well as behaviour, sex-specific phenotypes and reproductive maturation (Wilson *et al*., 1976). These genes share a common origin with the major royal jelly protein (MRJP) genes (Drapeau *et al*., 2006) which are crucial in caste development in honeybees. Like the MRJP genes, *yellow* genes in honeybees have diverse spatial and temporal expression patterns. As our focus in *A. plantaginis* has been on wing tissue, we are missing expression of genes in other tissues that could be linked to other traits. Thus, it is not impossible to imagine that a *yellow* gene could have a similar function to a MRJP in regulating the development of a complex phenotype. Recent work with *Bicyclus anynana* showed that *yellow* functions as a repressor of male courtship (Connahs *et al*., 2022). On the other hand, sex-specific behavioural phenotypes of *yellow* mutants in *Drosophila* were found to be due to pigmentation effects (Massey *et al*., 2019), so more evidence is needed to suggest a functional role for *yellow* genes outside of pigmentation.

The duplication of *yellow-e* and surrounding regions in the white morphs suggests that the yellow morph is the ancestral form. *Valkea* could have evolved in a stepwise fashion, first as a tandem duplication then with a stop codon mutation altering the gene structure. Gene duplications can facilitate adaptation and, in some examples, lead to polymorphism. The fact that the white allele is dominant also supports the hypothesis that yellow is ancestral. Such invasions of new adaptive alleles are facilitated when the new allele is dominant, as it is then also expressed when heterozygous, i.e. the Haldane’s sieve effect. Melanism, for example, has repeatedly evolved in mammals due to dominant and semidominant mutations in the *Mc1r* locus which have become fixed (Hoekstra, 2006).

*Valkea* could represent an example of neofunctionalization, where the duplicated gene gains a different function to the original gene copy. In the CRISPR mutants, both forewings and hindwings became yellow, and thus we suspect that *valkea* is controlling hindwing colour while *yellow-e*, which was likely also knocked out due to the similarity of the sequences, controls forewing colour. Those with the mutant phenotype showed only small deletions or insertions around the target site. By combining multiple guides we may expect to see larger deletions (Mazo-Vargas *et al*., 2022), however none of the eggs that were injected with more than one guide survived past the larval stage, suggesting that large deletions in *yellow* genes reduce fitness.

Contrary to previous work that attributed red and yellow colours to pterins in another tiger moth species (Gawne and Nijhout, 2019), we found high levels of 4-AHPEA in the yellow wings confirming the presence of pheomelanins. These pigments have been widely associated with red and yellow colours in mammals (e.g. Mcgraw and Wakamatsu, 2004), but only relatively recently described in insects and likely to be more widespread than previously thought. Yellow colours can also be produced by NBAD sclerotins which are sclerotizing precursor molecules made from dopamine and these have an important role in the sclerotization pathway for hardening the insect cuticle (Andersen, 2007; Barek, Evans and Sugumaran, 2017) before becoming involved in melanisation (Barek *et al*., 2018). Thus, we suggest that the yellow colour arises partly from the NBAD sclerotins and partly from the presence of pheomelanin pigments, which has been proposed in other Lepidoptera (Matsuoka and Monteiro, 2018). While some 4-AHPEA also occurred in white wings, this may be due to its role in production of cross-linking cuticular proteins and chitin during sclerotization (Sugumaran, 2010). Upregulation of genes on the white allele could be acting as a repressor of the generation of yellow colour. If we may speculate, *valkea* could impact the catalysis of dopamine, having cascading effects down the pathway resulting in the lack of yellow pigmentation. We suspect that *yellow* family genes play multiple roles within the melanin production pathway. In the wood tiger moth, *yellow* affects the conversion of DOPA into black dopamine melanin (Galarza, 2021). *Yellow-e* in particular has been linked to larval colouration in *Bombyx mori* (Ito *et al*., 2010) and adult colour in beetles (Wang *et al*., 2022), while another gene, *yellow-f*, has a role in eumelanin production (Barek *et al*., 2018).

In summary, we identified a structural variant which is only present in white morphs of *A. plantaginis*. This region contains a previously undescribed gene, *valkea*, which when knocked out results in yellow wings. The presence of a regulatory element controlling wing colour and other traits via multiple downstream effects could explain how multiple traits are linked to wing colouration. This complex polymorphism allows multiple beneficial phenotypes to be inherited together, whereas recombination would separate multiple loci leading to maladapted individuals. Our results provide the basis for further exploration of the genetic basis of covarying behavioural and life-history traits, and offer an intriguing example for the role of gene duplications in adaptive variation.

## Methods

### Sampling

Homozygous lines of white (WW) and yellow (yy) *Arctia plantaginis* moths were created from Finnish populations at the University of Jyväskylä, Finland. Larvae were fed with wild dandelion (*Taraxacum sp*.) and reared under natural light conditions, with an average day temperature of 25°C and night temperature between 15-20°C. For the crosses, a heterozygous male, created from crossing a heterozygous male with a homozygous yy female, was backcrossed with a yy female. This was repeated to obtain four families totalling 172 offspring and 8 parents (Table S5). Samples from wild populations were caught in Southern Finland (n=10) and Central Finland (n=20), where male morphs are either white or yellow, Estonia (n=4), where males are mostly white, and Scotland (n=4), where males are yellow (Table S6). In addition, we included 8 samples which are F1 offspring of wild Scottish samples. Forty pupae with known genotypes from lab populations (20 WW and 20 yy) were used for the RNA extractions.

### DNA extraction and sequencing

For the lab crosses, DNA was extracted from two legs crushed with sterilised PVC pestles using a Qiagen DNeasy Blood & Tissue kit, following the manufacturer’s instructions. Library preparation and GBS sequencing were performed by BGI Genomics on an Illumina HiSeq X Ten. For the wild samples, DNA was extracted from the thoraces also with a Qiagen kit. Library preparation and sequencing were performed by Novogene (Hong Kong, China). 150bp paired-end reads were sequenced on an Illumina NovaSeq 6000 platform.

### Linkage mapping analysis

FASTQ reads were mapped using bowtie v2.3.2 (Langmead and Salzberg, 2012) to the yellow *Arctia plantaginis* scaffold-level genome assembly (Yen *et al*., 2020). BAM files were sorted and indexed using SAMtools v.1.9 (Li *et al*., 2009) and duplicates removed using PicardTools MarkDuplicates (broadinstitute.github.io/picard). Twelve samples which had aligned <30% were removed. Reads of the remaining samples had an average alignment of 94%. SNPs were called using SAMtools mpileup with minimum mapping quality set to 20 and bcftools call function. Lep-MAP3 (Rastas, 2017) was used for linkage map construction and we ran the following modules: ParentCall2 which called 105,622 markers, Filtering2, SeparateChromosomes2 with lodLimit=5 and sizeLimit=100, JoinSingles2All and OrderMarkers2 with recombination2=0 to denote the lack of female recombination. Genotypes were phased using the map2genotypes.awk script included with Lep-MAP3. Markers were named based on the genomic positions of the SNPs in the reference genome and the map.awk script, and this was used to further order the markers within the linkage groups. This resolved 30 linkage groups. Although we expect that there are 31 chromosomes in the moth genome, we suspect that the sex chromosome is missing in this assembly as the yy individual used in the genome assembly was female (Yen *et al*., 2020). A small number of markers which caused long gaps at the beginning or end of linkage groups were manually removed, leaving the final map 948.7cM long with 19,803 markers. Markers were well distributed so we began the first analyses with this map. A linkage map was also assembled using sequences aligned to the white reference and this separated into 31 linkage groups.

The QTL analysis was carried out in R/qtl (Broman *et al*., 2003). Genotype probabilities were calculated before running a genome scan using the *scanone* function with the Haley-Knott method and binary model parameters, and including family as an additive covariate. The phenotype was labelled as either 0 (Wy) or 1 (yy). We ran 5000 permutations to determine the significance level for the QTL LOD scores. The *bayesint* function calculated the 95% Bayesian confidence intervals around the significant marker.

### Analysis of whole genome sequences

FASTQ reads were mapped to the yellow *A. plantaginis* genome assembly (Yen *et al*., 2020) using BWA-MEM v7.17 (Li, 2013). As before, BAM files were sorted and indexed, and duplicates were removed. Genotyping and variant calling was carried out with the Genome Analysis Toolkit (GATK) (McKenna *et al*., 2010). Variants were called using HaplotypeCaller (v.3.7) in GVCF mode, then gVCFs combined with GenomicsDBImport (v.4.0). Joint genotyping was run with GenotypeGVCFs, set with a heterozygosity of 0.01, and SNPs were called using SelectVariants. Finally, the set of 20,787,772 raw SNPs were filtered using VariantFiltration and thresholds: quality by depth (QD > 2.0), root mean square mapping quality (MQ > 50.0), mapping quality rank sum test (MQRankSum > −12.5), read position rank sum test (ReadPosRankSum > −8.0), Fisher strand bias (FS < 60.0), and strand odds ratio (SOR < 3.0). A set of 5,227,288 SNPs passed the filtering.

We carried out a Genome Wide Association Study (GWAS) using the R package GenABEL v.1.8 (Aulchenko *et al*., 2007). The set of filtered SNPs were converted to BED format with PLINK2, keeping only biallelic SNPs (cog-genomics.org/plink/2.0/). Sites which were not in Hardy-Weinberg equilibrium (p<0.01), or had a call rate of <0.5, were excluded. Following this, 381,266 sites were retained across 40 individuals (out of 57). To account for population stratification, we performed multidimensional scaling on kinship and identity-by-state (IBS) information estimated from the data, and included this as a covariate in the association test. Significance levels were calculated using Bonferroni corrected thresholds to account for multiple testing. Central and Southern Finnish populations were pooled for this analysis, based on a previous principal component analysis of these samples (Yen *et al*., 2020).

In Yen et al. (2020), many of these samples were processed in the same way but aligned to the white genome assembly (based on a white individual). Read depth of the W-mapped samples was calculated in 1kb windows across the candidate region using BEDtools (v.2.20.1) multicov (Quinlan and Hall, 2010). For visualisation, lines were smoothed using LOESS and span=0.01 within ggplot2.

For analysis of structural variants, sequences from the white and yellow genome assemblies were aligned using MAFFT v7.450 (Katoh and Standley, 2013) and viewed with Geneious. Our focal sequence, scaffold 419 in the white genome, is the reverse complement of scaffold 206 in the yellow genome.

### Identification of candidate genes and tree construction

To identify candidate genes in the QTL interval and GWAS region, we ran a protein BLASTP v.2.4.0 search to identify *Heliconius melpomene* (Hmel2.5) proteins homologous to predicted *A. plantaginis* proteins in the region from the genome annotation. Informative gene names were obtained by performing a BLASTP search with the *H. melpomene* proteins against all *Drosophila melanogaster* proteins in FlyBase v.FB2020_01 (flybase.org/blast).

For the *yellow* gene tree, we used Lepbase (Challi *et al*., 2016) to search for *yellow* genes in *Bombyx mori* (ASM15162v1). We identified *yellow-e* in *H. melpomene* by searching for major royal jelly proteins, then comparing protein sequences of these against *Drosophila* proteins in Flybase. The sequence for *Dmel yellow-e* was downloaded from Flybase. To make the tree, coding sequences of all genes were aligned in Geneious using MAFFT v7.450 (Katoh and Standley, 2013), then the tree was constructed with PhyML using 10 bootstraps (Guindon *et al*., 2010).

### Differential gene expression

We dissected the wings out of the pupae in Cambridge, UK. Pupae and larvae were sent to Cambridge from Jyväskylä and were kept between 22 and 30°C. Pupae were sexed and only males were used. Dissections were made at 4 different stages: 72 hours after pupation (72h), 5 days after pupation (5d; counting 0-24 first hours after pupation as day 1), pre melanin deposition (Pre-mel) and post melanin deposition (Mel). We sampled 5 individuals per stage and genotype. Hindwings and forewings were stored separately in RNA-later (Sigma-Aldrich) at 4 °C for 2 weeks and later transferred to −20 °C, while the rest of the body was stored in pure ethanol. Only hindwings were used for RNAseq analysis.

Total RNA was extracted from hindwing tissue using a standard hybrid protocol. First, we transferred the wing tissue into Trizol Reagent (Invitrogen) and homogenised it using dounce tissue grinders (Sigma-Aldrich). Then, we performed a chloroform phase extraction, followed by DNase treatment (Ambion) for 30 mins at 37°C. We measured the concentration of total RNA using Qubit Fluorometric Quantitation (Thermofisher) and performed a quality check using an Agilent 4200 TapeStation (Agilent). The extracted total RNA was stored at −20°C before being sent to Novogene UK for sequencing. Each individual was sequenced separately, with a total of 40 individual samples sequenced (5 individuals per stage and genotype).

We performed quality control and low-quality base and adapter trimming of the sequence data using *TrimGalore!* We then mapped the trimmed reads to the two *A. plantaginis* genomes using STAR (Dobin *et al*., 2013). We performed a second round of mapping (2pass) including as input the output splice junctions from the first round. The *A. plantaginis* genome annotations WW and YY were included in each round of mapping respectively. We then used *FeatureCounts* to count the mapped reads. Finally, we used DESeq2 to analyse the counts and perform the DE analysis.

To identify the gene or genes controlling the development of wing colour in *A. plantaginis*, we performed a genome wide differential expression analysis using limma-voom (Ritchie *et al*., 2015). First, we defined a categorical variable, ‘GenStage’, with 8 levels containing the genotype and stage information of every individual sample (e.g. YY72h, WW72h, YY5days, etc.). Then, we built the design matrix fitting a model with GenStage as the only fixed effect factor contributing to the variance in gene expression and included family as a random effect factor (gene expression ∼ 0 + GenStage + (1|Family)). We then filtered lowly expressed genes using the filterByExpr function in limma, which resulted in a reduction of the number of tested genes from 17,615 to 11,330 genes in the Y-mapped analysis and 17,930 to 10,920 in the W-mapped one. Then, we normalised the expression of the genes using the calcNormFactors function with TMM normalisation in limma and fit the design matrix using the voom function. We built a contrast matrix including the comparisons of interest, in which we compared the expression of the genotypes in each stage (i.e. h72=WW72h-YY72h, d5=WWd5-YYd5, Premel=WWPremel-YYPremel, Mel=WWMel-YYMel), and fit the contrast matrix to the data using the contrasts.fit function. Finally, we used the eBayer function on the fit dataset and we extracted the list of genes that are differentially expressed in each stage using the Benjamini-Hochberg procedure to correct for multiple testing. We evaluated the genome wide gene expression using multidimensional scaling (MDS) using the plotMDS function of the limma package.

### Orthology assignment

To infer genome wide orthology between *A. plantaginis* and *D. melanogaster*, we used OrthoFinder (v2.5.4) (Emms and Kelly, 2019). We used proteomes from six Lepidoptera species, *Plutela xylostella* (GCA_905116875_2), *Bombyx mori* (GCF_014905235_1), *Spodoptera frugiperda* (GCF_011064685_1), *Parnassius apollo* (GCA_907164705_1), *Pieris macdunnoughi* (GCA_905332375_1) and *Pararge aegeria* (GCF_905163445_1) and *Drosophila melanogaster* (GCF_000001215_4). We ran the primary_transcript.py utility from OrthoFinder to extract only one transcript per protein, and then ran OrthoFinder with default settings.

### CRISPR/Cas9 genome editing

Guide RNAs were designed within the first 3 exons of *valkea* in the white genome annotation using Geneious (v. 2022.1.1). Guides were chosen with minimal off-target effects, high activity scores and high specificity scores based on the Geneious algorithm (Table S7). Guides in the first two exons of *valkea* showed off-target sites in *yellow-e*, however those in exon 3 showed no off-target sites. Guide RNAs were synthesised by Sigma-Aldrich. Moths from the greenhouse populations, originating from Finnish and Estonian populations, at the University of Helsinki, Finland, were genotyped using DNA extracted from leg tissue using the Chemagic DNA tissue kit (Chemagen) and the primers detailed below. They were paired and left to mate overnight. Females were watched over the next 3-4 days for egg laying, and the eggs were removed and injected less than 6 hours after laying. Eggs were glued to microscope slides and injected with a 1:1 sgRNA/Cas9 mix with phenol red dye using pulled borosilicate glass capillaries. The injection mix contained 1ug/ul Cas9, 500ng/ul sgRNA and 0.5% phenol red. Guides and Cas9 were diluted using low concentration TE buffer. Different combinations of guides were also injected in some samples, in which case these were mixed in a 1:1 ratio. Injections were performed using a MPPI-3 pressure injector with back pressure unit (ASI). In total, we injected 1223 eggs from 18 [WW x WW] or [WW x Wy] crosses. Larvae were kept individually in petri dishes and fed daily with dandelion leaves. After eclosion, legs were taken from adults for DNA extraction. Library preparation and whole genome sequencing (using Illumina NovaSeq 6000) of 5 CRISPR mutants was performed by CeGaT (Tübingen, Germany).

### Genotyping white and yellow alleles

We used Primer3 to design primers within the duplicated region. Primers were expected to only amplify in WW and Wy individuals. The alignment of the white and yellow sequences was then used to design primers for genotyping the locus (Table S2). We looked for short insertions or deletions that were fixed between the WW and yy within the *valkea/yellow-e* region, and put primers around these structural variants. Primers were tested on DNA extractions from moths of known genotypes, including both sexes, wild and lab samples. We used Sanger sequencing of the PCR product to confirm the correct sequences were amplified. A set of primers successfully amplified a 449bp region downstream of *valkea* within the duplication. This amplified in WW and Wy samples, but not in yy (Appendix Figure 1). We found that white alleles have a 35bp deletion within an intron of the *yellow-e* gene. We amplified a 163bp region around this (YY_tarseq_206_arrow:9,846,212-9,846,375) using a standard PCR protocol which allowed us to identify the allele based on the size of the PCR product. Yellow alleles produce the full 163bp sequence, while white alleles produce a smaller 128bp product (Appendix Figure 1). Heterozygotes have a copy of each and show both bands on a gel.

### Pigment analysis

#### Solubility and fluorescence tests

Five hindwings from each morph were placed in two separate solvents (0.1M NaOH and 90% MeOH) and left overnight. The supernatant was analysed with an Agilent Cary 8454 UV-Visible spectrophotometer and the spectra compared to known spectra for various pigments. A UV lamp (Philips TL8W/08 F8T5/BLB) was used to test for fluorescence on the wings. The presence of carotenoids was tested by placing wings into 1ml of pyridine and leaving at 95°C for 4 hours (McGraw *et al*., 2002). To these we added 1ml of 1:1 hexane:tert-butyl methyl ether and 2ml of water before shaking and leaving overnight. Again, the supernatant was measured with the spectrometer.

#### High-Performance Liquid Chromatography (HPLC) to test to for eumelanin and ommochrome pigments

To determine the type of melanin producing the black colour on the wings, we cut out approximately 5mg of the black sections of the wings, from both females and males. Eumelanin analysis was carried out according to (Borges *et al*., 2001). Each sample was added to a tube containing 820μl 0.5 M NaOH, 80μl 3% H_2_O_2_ and an internal standard (48nmol phthalic acid) and heated in a boiling water bath for 20 minutes. Once cool, 20μl of 10% Na_2_SO_3_ and 250μl of 6M HCl were added. Samples were then extracted twice with 7ml of ethyl acetate. Ethyl acetate was dried at 45 °C under a stream of nitrogen. The residue was dissolved into 0.5ml of 0.1% formic acid.

We carried out high-performance liquid chromatography on an Agilent 1100 HPLC. 20μl of the sample was injected into a Waters Atlantis T3, 100 × 3.0mm i.d. analytical column (Waters, Milford, MA, USA). The column was set to 25°C and analytes were detected at wavelength 280nm. The HPLC mobile phase consisted of two eluents: UHQ-water/MeOH (98/2; v/v) with 0.1% formic acid and UHQ-water/MeOH (40/60; v/v) with 0.1% formic acid. Flow rate was 0.4 ml/min and the used gradient started with 100% of eluent A and ramped evenly from time 0 to 15 minutes to 40:60 (A:B; v/v), held at 40:60 for 6 minutes and ramped evenly back to initial eluent composition (100% A) over 5 minutes. We compared chromatograms obtained from the samples to the chromatograms obtained from synthetic melanin, ink from sepia officinalis and black human hair.

HPLC was also applied to observe the possible presence of ommochrome pigments. Injection volume was 10µl and for the separation we used the same Waters Atlantis T3 column (100 × 3.0 mm i.d.) set to 30°C. Solvent A was UHQ-water and B was acetonitrile (ACN), both containing 0.1% formic acid. Flow rate was 0.4ml/min and the used gradient was as follows: Initial flow ratio was 98/2 water/ACN (v/v) ramping then evenly from time 1 to 15 min to 30:70 water:ACN (v/v), held for 1.5 min and then ramped evenly back to initial eluent composition over 0.5 minutes. The column was stabilised for 7 minutes before a new run.

#### Pheomelanin analysis

Samples were analysed for pheomelanin content according to the method of Kolb *et al*. (1997) with modifications. A 2mg sample was placed in a screw-capped tube with 100μl water, 500μl ∼55-58% hydrogen iodide (HI), and 20μl 50% hypophosphorous acid (H_3_PO_2_). Samples were capped tightly and hydrolysed for 20 hours at 130°. After cooling, samples were evaporated under nitrogen flow, then dissolved in 1ml of 0.1M HCl and purified with solid phase extraction. Strata SCX cartridges were preconditioned with 2ml of methanol, 3ml of water and 1ml of 0.1M HCl. Sample was then applied to the cartridge, washed with 1ml of 0.1M HCl, and finally eluted with 1ml of methanol (MeOH): 0.5M ammonium acetate (NH_4_CH_3_CO_2_) (20:80 v/v).

Hydrogen Iodide hydrolysis products were determined by a Dionex high-performance liquid chromatograph equipped with pulsed amperometric detection (HPLC/PAD). A Phenomenex Kinetex C18 column (150 × 4.6 mm i.d.; 5µm particle size) with a gradient elution (Table S8) at a flow rate of 0.9 ml min^-1^ with the eluents: (A) sodium citrate buffer (Hines *et al*., 2017) in ultra-high-quality water (internal resistance ≥ 18.2 MΩ cm; Milli-Q Plus; Millipore, Bedford, MA, USA) and (B) methanol were used for the separation. Dionex ED-50 pulsed amperometric detector (Dionex, Sunnyvale, CA, USA) equipped with a disposable working electrode by using a Dionex waveform A with potentials presented in Table S9 was used for detection. The preparation method for the 4-AHP, 3-AHPEA and 4-AHPEA standards used in calibration is described in (Wakamatsu *et al*., 2014).

## Acknowledgements

We thank Alma Oksanen, Kaisa Suisto and the greenhouse staff for insect rearing, and Elisa Salmivirta and Sari Viinikainen for lab assistance. Thanks to Emeritus Prof. Shosuke Ito for kindly providing pheomelanin standards, and to Bodo Wilts for advice on the pigment analyses. We thank Muktai Kuwalekar, Claudius Kratochwil, Rachel Blow, Ian Warren, Tom Generalovic and Joe Hanly for providing equipment and advice regarding the CRISPR injections.

## Data availability

Scripts and data for the QTL, GWAS and DE analyses can be found at doi:10.5281/zenodo.6560628. RADseq, RNAseq data, and WGS of CRISPR samples were deposited to SRA under study accession number PRJNA937225. Raw sequencing data of wild samples has previously been deposited in ENA, study accession No. PRJEB36595.

