## supplement for "Colour polymorphism associated with a gene duplication in male wood tiger moths"

### Figure supplements

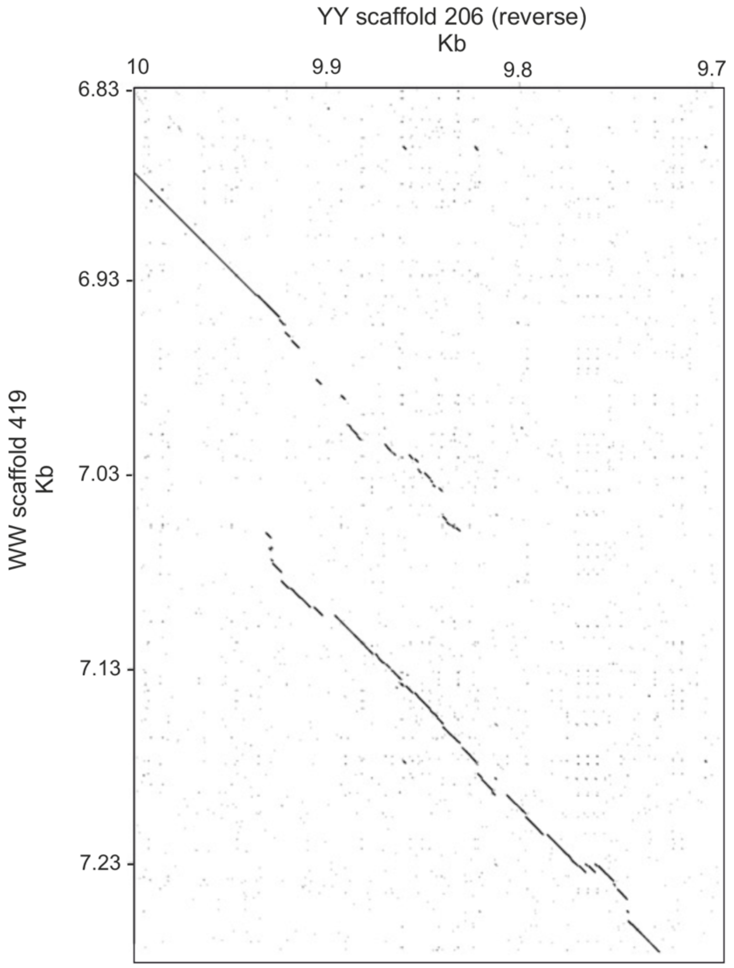


Figure 2 – figure supplement 1. Dotplot of the alignment between a region of scaffold 419 from the white reference and scaffold 206 from the yellow reference.

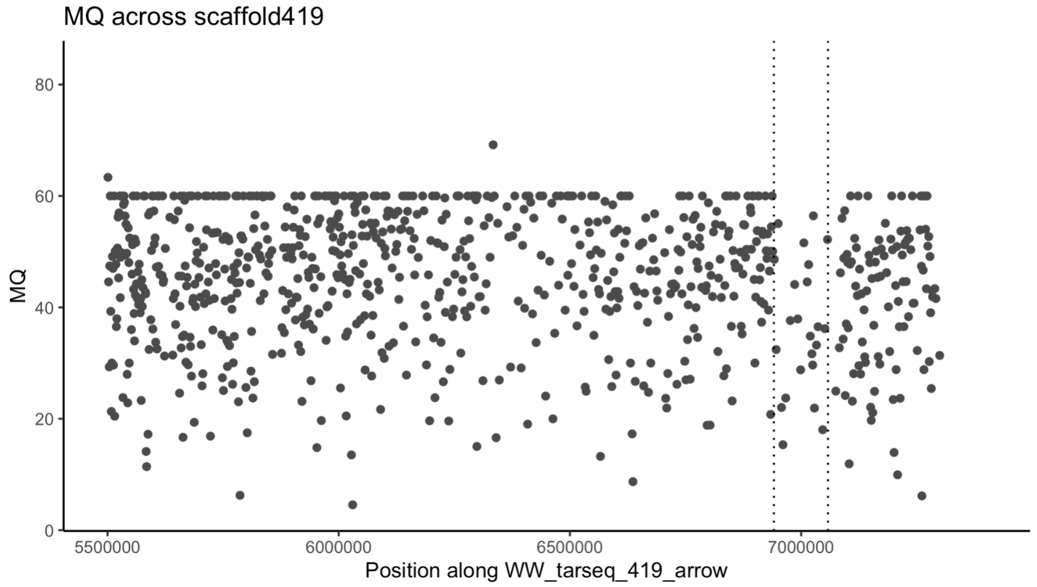


Figure 2 – figure supplement 2: Mapping quality (MQ) across the latter half of scaffold WW_tarseq_419_arrow. MQ is lower within the duplicated region (between the dotted lines): average MQ for the duplication including all samples is 36.52, whereas for the rest of the scaffold, not including the duplication, it is 44.16.

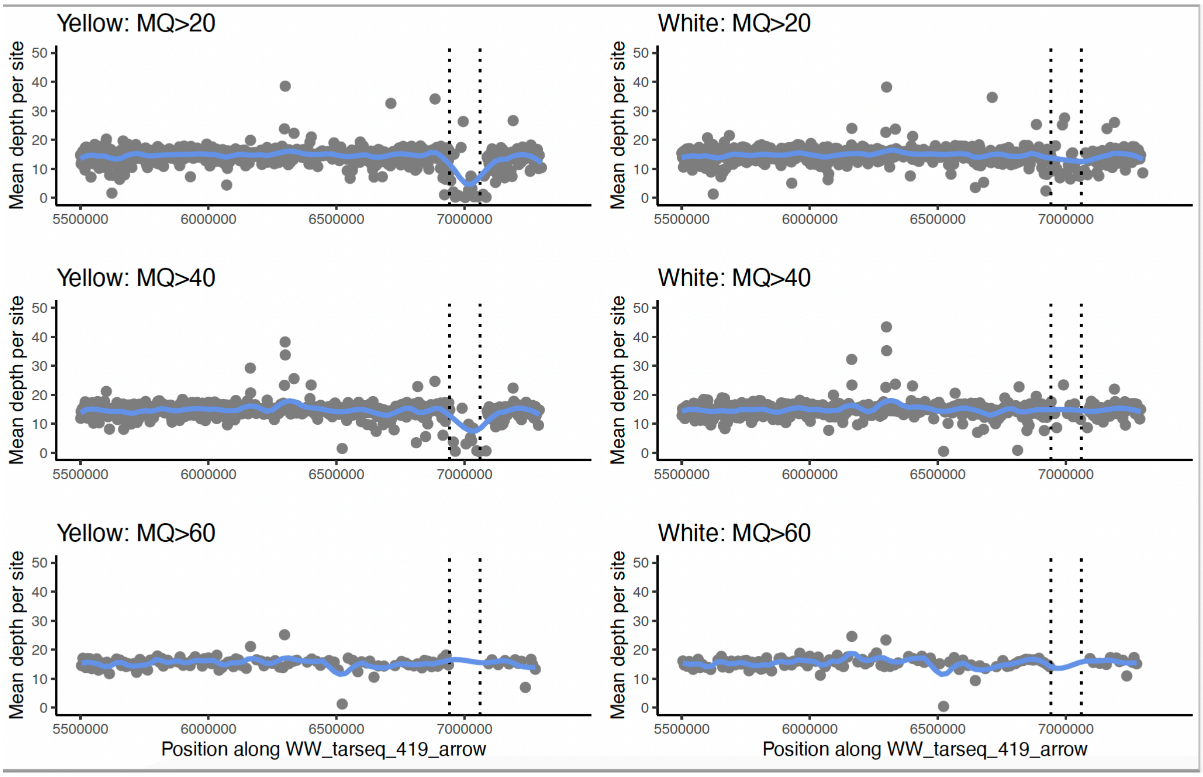


Figure 2 – figure supplement 3: When filtering for increasing mapping quality, more reads are lost within the duplication in yellow samples compared to white samples, suggesting that reads in this region are more likely to be mismapped than in white samples.

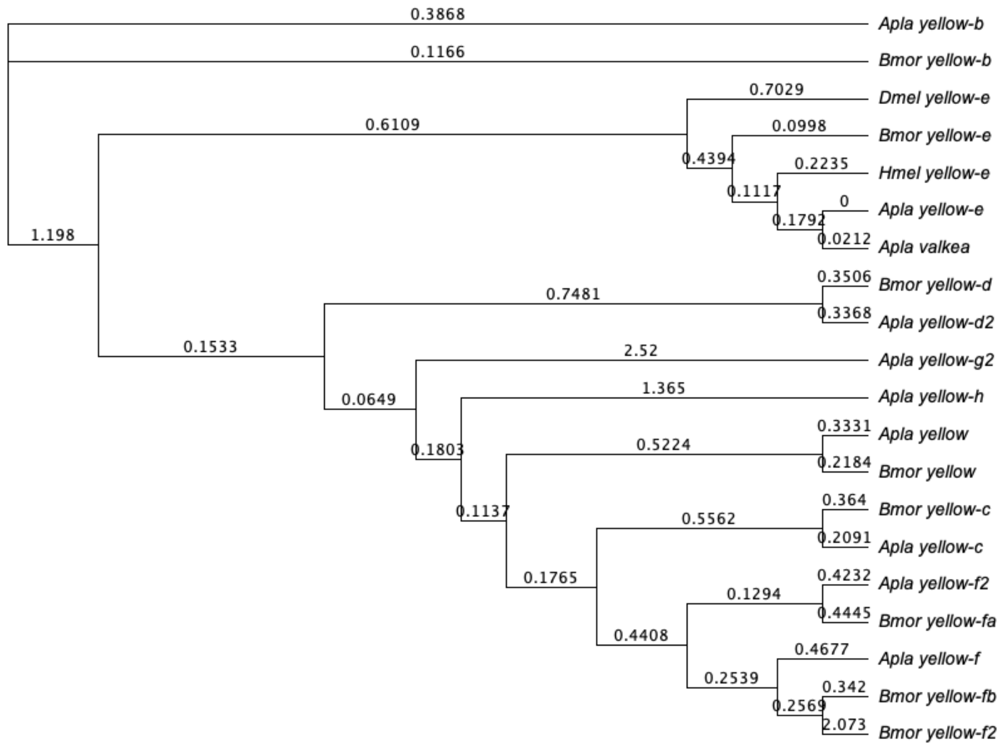


Figure 2 – figure supplement 4: Tree of coding sequences for all *yellow* family genes found in *A. plantaginis* and *B. mori. Yellow-e* sequences from *D. melanogaster* and *H. melpomene* are also included. Values are substitutions per site.

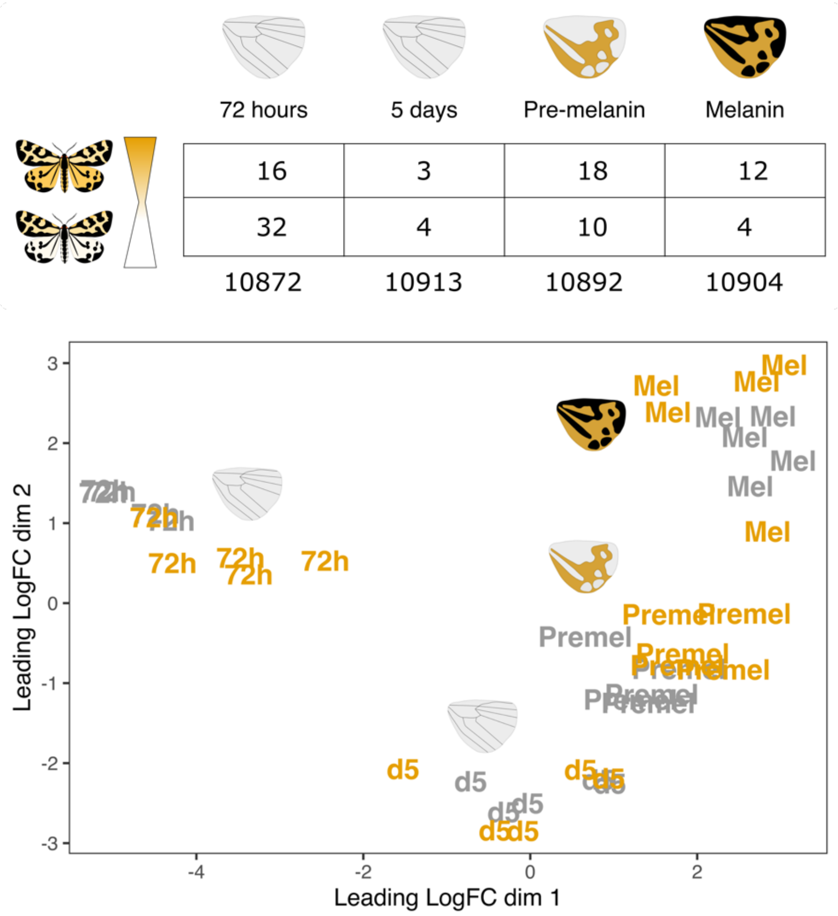


Figure 3 – figure supplement 1: Genome wide expression patterns are shaped by developmental stage. Multidimensional scaling of gene expression shows that samples cluster by developmental stage rather than male colour morph. Developmental stage names have been abbreviated: 72h=72 hours, 5d=5 days, Premel=pre-melanin and Mel=melanin.

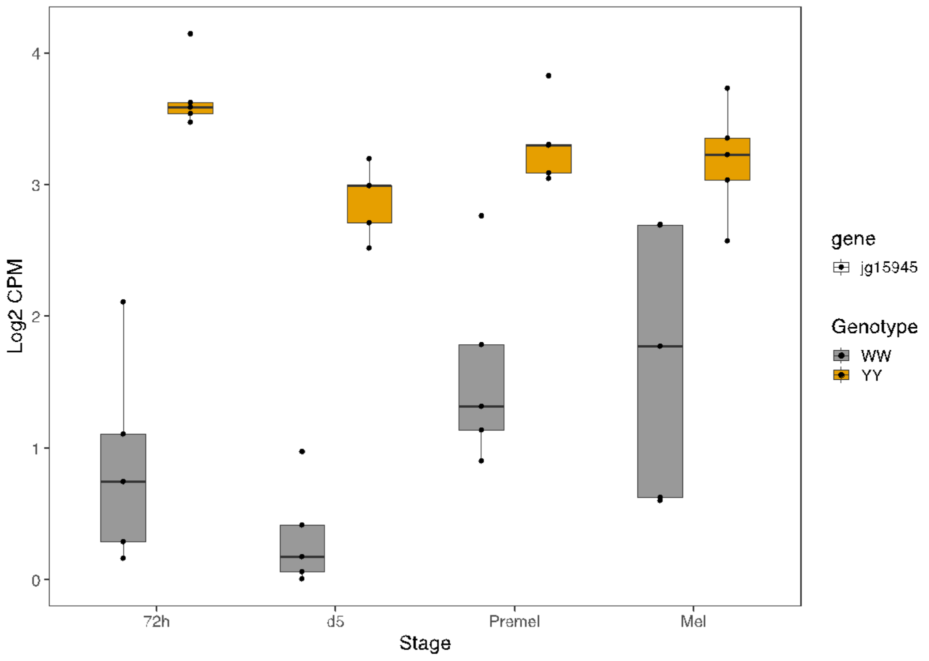


Figure 3 – figure supplement 2: Expression of the gene 'jg15945' across the 4 developmental stages.

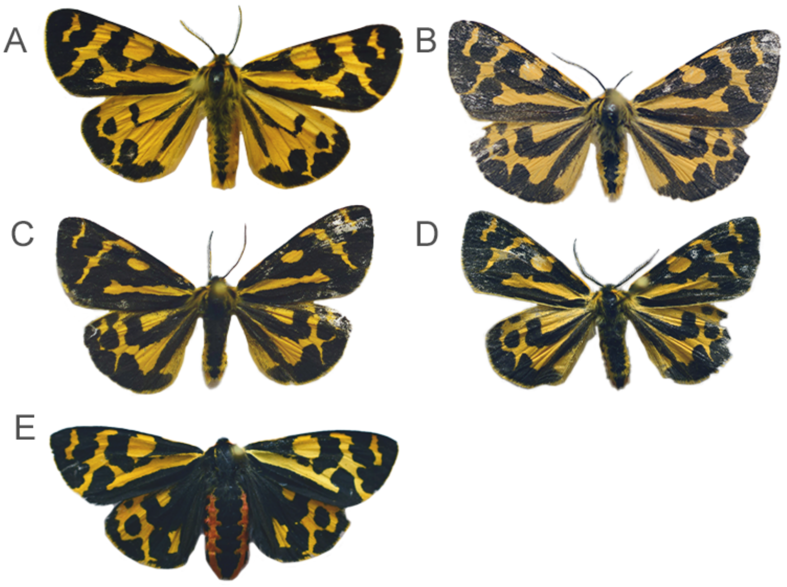


Figure 4 – supplement 1: The four males showing the mutant phenotype. Variation in melanin amount is likely due to the lineage, with (A) coming from the Estonian line, and (B-D) coming from the Finnish line. (E) Female showing a mosaic knockout phenotype with left forewing showing similar colour to the male mutants.

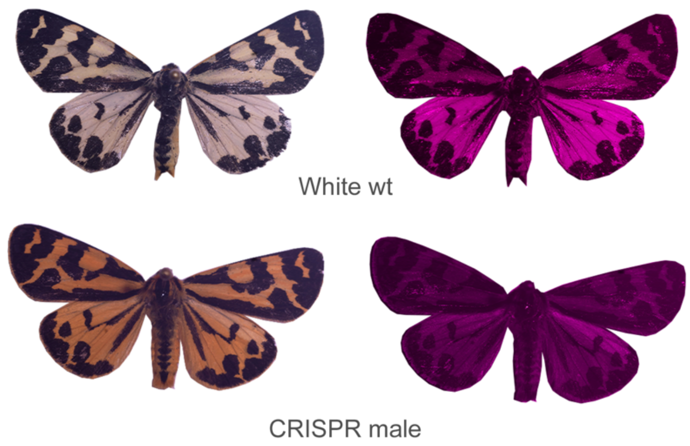


Figure 4 – supplement 2: Wildtype WW morphs show UV reflectance on the dorsal sides of the wings. This is not seen in the CRISPR knockout males.

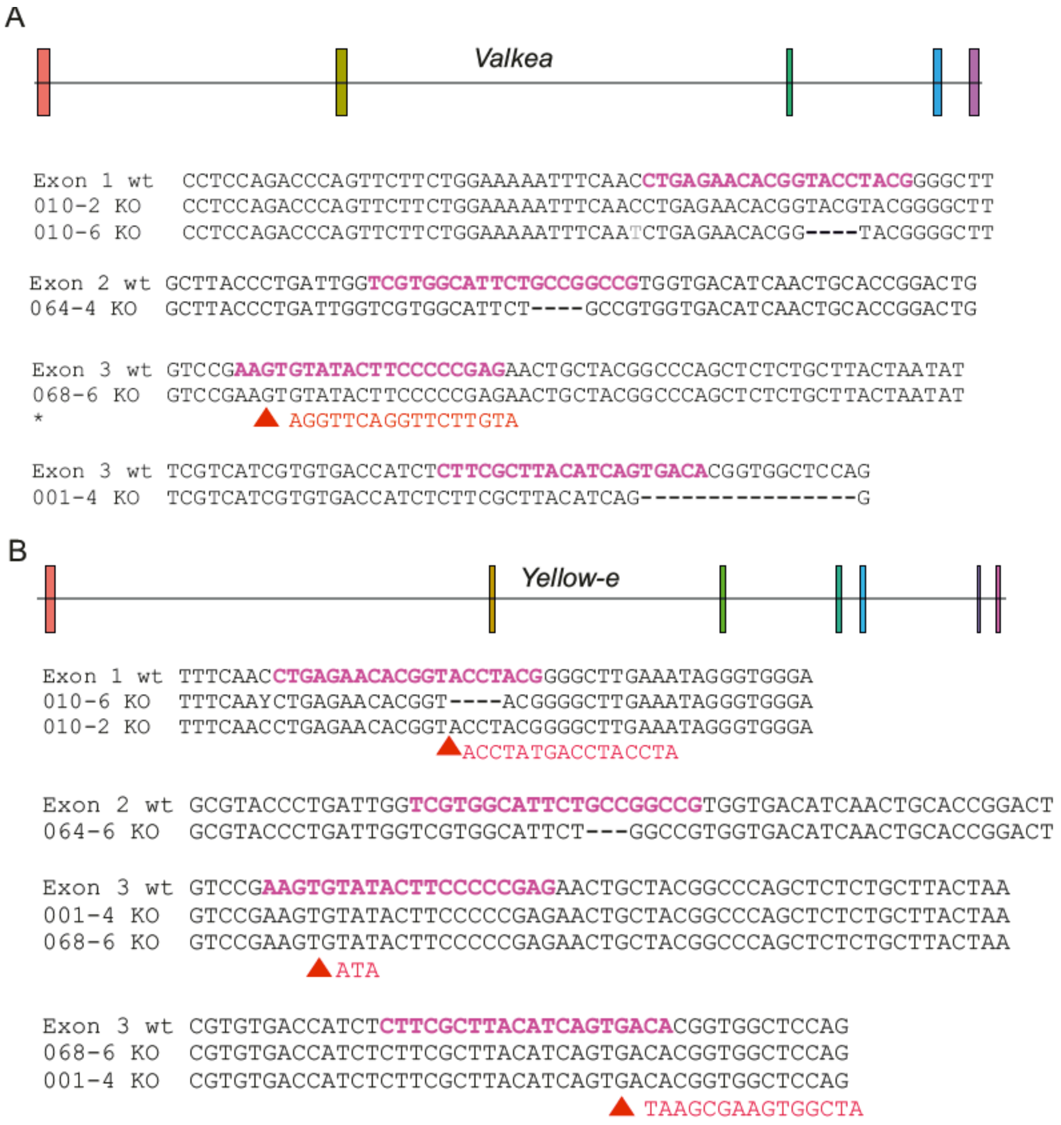


Figure 4 – figure supplement 3: CRISPR knockouts confirmed using sequence data of the five individuals with visible changes in phenotype. The top row of each block shows the reference sequence, and the rows underneath are the mutant individuals. Pink sequences denote the guide RNA sequence. Four different guides produced mutants (one in each of exons 1 and 2, and two in the third exon). Red arrows show insertions. (A) Three out of five samples show small deletions in *valkea* close to the target sites. One sample had an insertion. (B) In the corresponding sites in *yellow-e,* two samples showed evidence of deletions and three had insertions.

**Appendix figures**

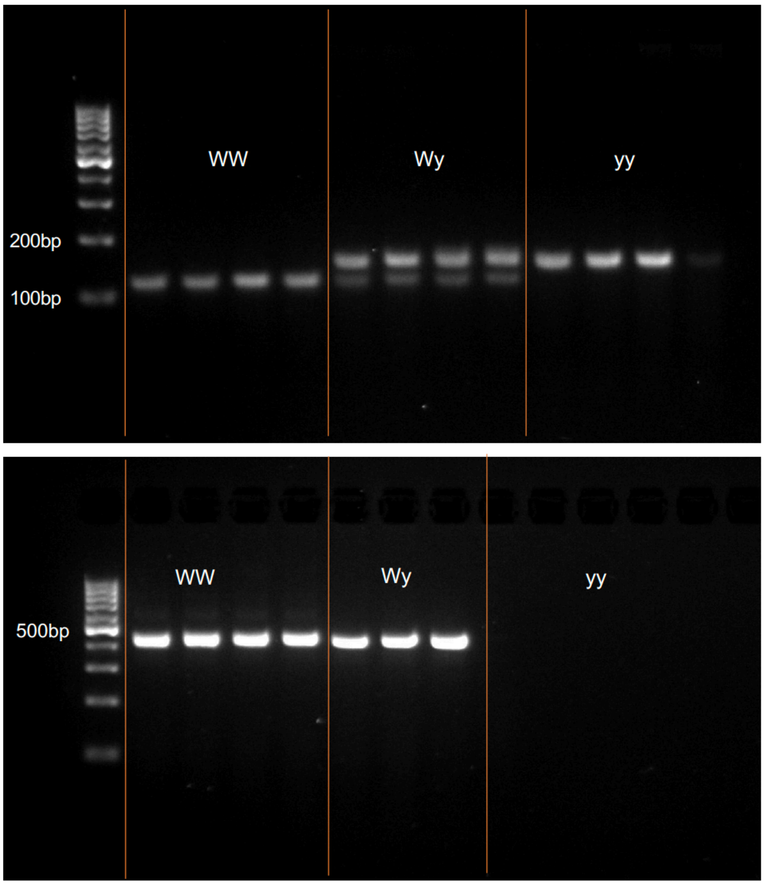


Appendix Figure 1: Example gel images for the genotyping primers. (A) Yellow alleles have the larger 163bp band, while white alleles have a 35bp deletion producing the smaller ~128bp band. Heterozygotes have a copy of each. (B) A 449bp region within the duplication amplifies only in samples with 1 or 2 white alleles.

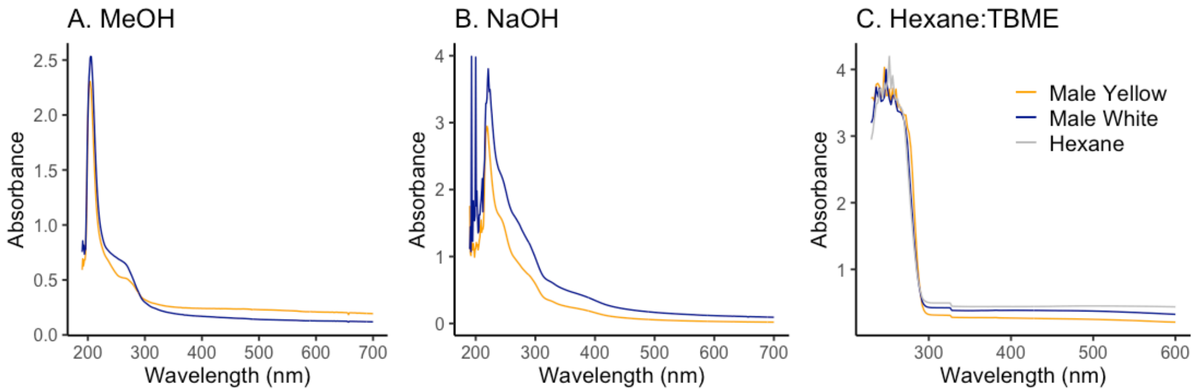


Appendix Figure 2: Absorbance curves for yellow and white male wings and female wings left in (A) methanol, which absorbs at 205nm, (B) sodium hydroxide, which absorbs in the UV range, and (C) hexane:tert-butyl methyl ether which absorbs between 195-210nm, as shown in the control measurement. Peaks outside of these values would suggest the presence of pigment compounds in the solvent, but these are not seen.

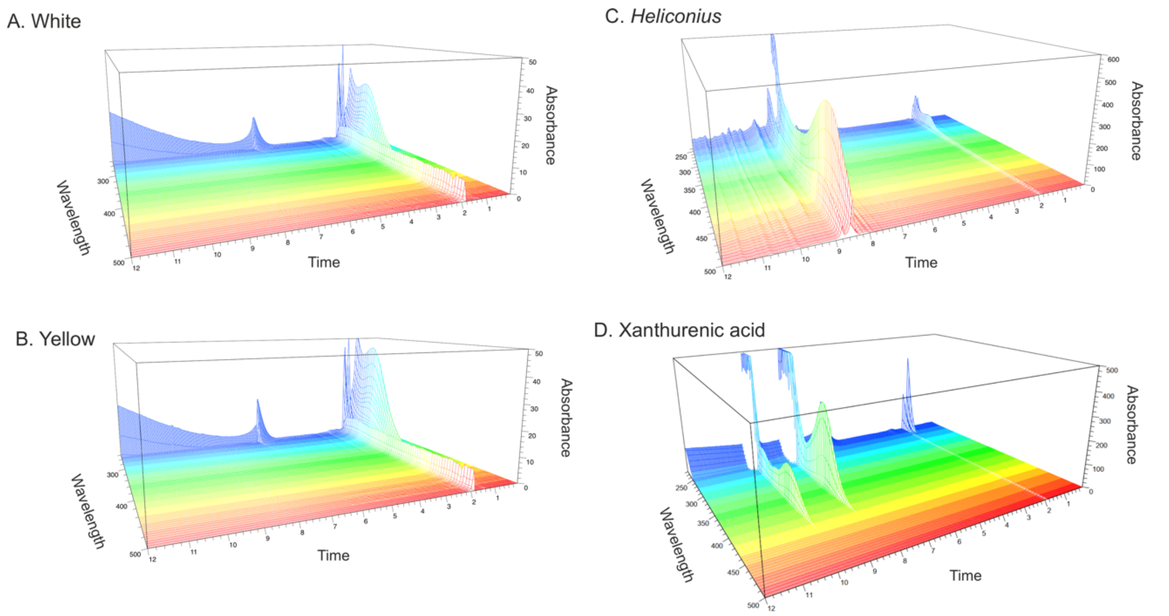


Appendix Figure 3: Spectral and chromatogram data obtained using HPLC to test for the presence of ommochrome pigments. White (A) and yellow (B) *A. plantaginis* morphs are compared to ommochrome-containing *Heliconius* wings (C) and a Xanthurenic acid standard (D).

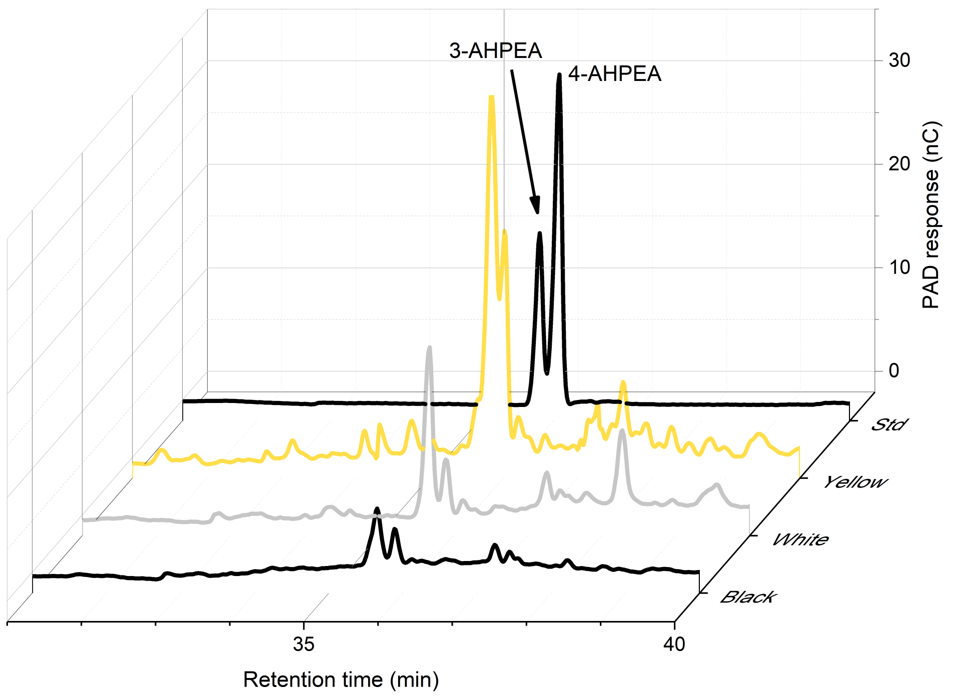


Appendix Figure 4. HPLC analysis for yellow, white, and black portions of the hindwing, plus the standard (Std). The highest levels of 4-AHPEA, a breakdown product of pheomelanin, is seen in the yellow wings.

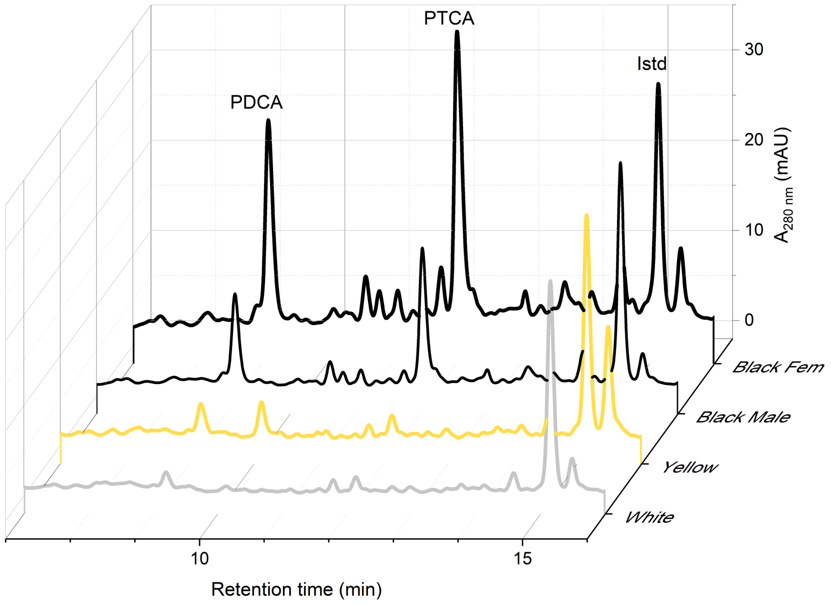


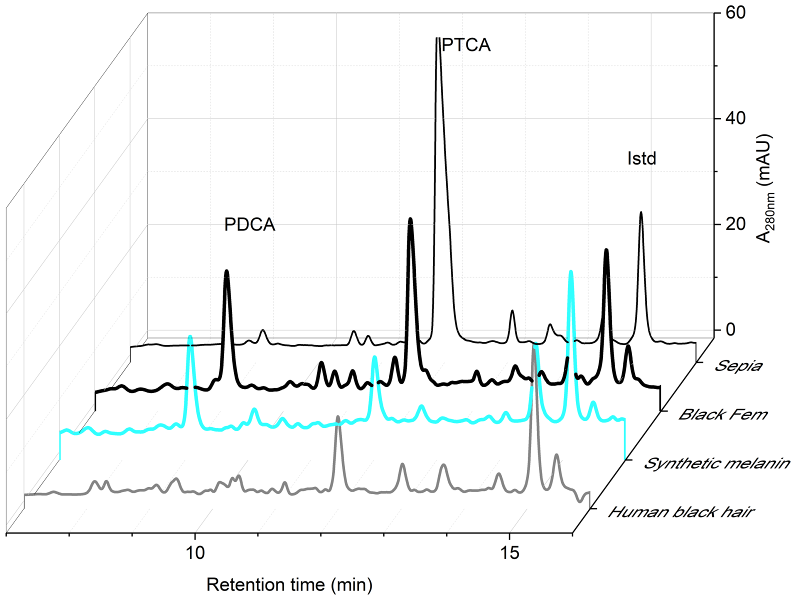


Appendix Figure 5: HPLC result for eumelanin analysis. The black portions of the wings show the presence of dopamine-derived eumelanin components (PDCA and PTCA). Phthalic acid is the internal standard (Istd). Below are results of eumelanin-containing samples for comparison – sepia, synthetic material and human hair.

**Supplementary tables**

Table S1: 21 genes are within in the QTL interval. Start and end positions shown are on scaffold 206 in the yellow *A. plantaginis* reference genome. Gene sequences were blasted against *Heliconius melpomene* and searched for in Flybase. Apla gene names from annotations produced by Yen et al. (2020).

| **Apla gene name (yellow ref)** | **Hmel gene hit** | **% identical** | **Hmel chr** | **Dmel gene hit** | **Blast score** | **E value** |
|---|---|---|---|---|---|---|
| jg6722 | HMEL032761g1.t1 | 45.8 | 15 | kis-PF | 29.261 | 3.32994 |
| jg6723 | unknown | | | | | |
| jg6724 | unknown | | | | | |
| jg6725 | HMEL032764g1.t1 | 80.9 | 15 | CG9853-PC | 234.958 | 7.86E-62 |
| jg6726 | HMEL004965g1.t1 | 68.7 | 15 | CG8128-PB | 240.736 | 9.40E-64 |
| jg6727 | HMEL004965g1.t1 | 61.2 | 15 | CG8128-PB | 240.736 | 9.40E-64 |
| jg6728 | HMEL004964g1.t1 | 43.6 | 15 | CG6262-PA | 29.261 | 0.764596 |
| jg6729 | unknown | | | | | |
| jg6730 | unknown | | | | | |
| jg6731 | HMEL004962g1.t1 | 81.4 | 15 | dimm-PB | 105.145 | 4.56E-23 |
| jg6732 | HMEL016957g1.t1 | 62.4 | 19 | CkIalpha-PG | 512.301 | 2.35E-145 |
| jg6733 | unknown | | | | | |
| jg6734 | HMEL004960g1.t1 | 69.7 | 15 | CG7322-PC | 184.882 | 5.40E-47 |
| jg6735 | HMEL032767g1.t1 | 85.2 | 15 | CG7322-PC | 209.92 | 1.57E-54 |
| jg6736 | HMEL010407g1.t1 | 75 | 15 | CG10311-PB | 142.51 | 1.76E-34 |
| jg6737 | HMEL032770g1.t1 | 59.7 | 15 | CG34307-PB | 39.2762 | 0.00975671 |
| jg6738 | HMEL010409g1.t1 | 72.1 | 15 | yellow h | | |
| jg6739 | HMEL010409g1.t1 | 46 | 15 | yellow h | 112.464 | 1.74E-25 |
| jg6740 | HMEL032771g1.t1 | 76 | 15 | CG8401-PB | 35.8094 | 0.0810412 |
| jg6741 | HMEL002092g1.t1 | 68.4 | 15 | yellow d2 | 290.426 | 2.27E-78 |
| jg6742 | HMEL032773g1.t1 | 86.7 | 15 | yellow e | 387.882 | 8.79E-108 |

Table S2: Primers used for genotyping. Tested using GoTaq Flexi buffer and GoTaq DNA polymerase, with annealing temperature of 57$^{o}$C for 35 cycles. ‘Ye12’ primers surround a small deletion in white alleles, producing a 163bp product from Y alleles and 128bp product from W alleles. ‘Dup5’ primers amplify a 449bp sequence within the duplicated sequence only in moths with at least one W allele. See Appendix figure 1 for gel images.

| **Primer name** | **Primer sequence** | **Position in YY genome** | **Position in WW genome** |
|---|---|---|---|
| ye12_F | ATGGCCGATTACGTCTTACGAC | YY_tarseq_206_arrow:9846212 | WW_tarseq_419_arrow:7150189 |
| ye12_R | CAACTAAATACAAAGTAGCTTCCCT | YY_tarseq_206_arrow:9846375 | WW_tarseq_419_arrow:7150353 |
| dup5C_f | ACTGACGTTTGTTTTGTCCCAA | NA | WW_tarseq_419_arrow:7052355 |
| dup5D_R | GGTGTGCATATTCCTGCTGT | NA | WW_tarseq_419_arrow:7052785 |

Table S3: List of differentially expressed genes found in the linkage group containing scaffold 419 (WW reference).

| *A. pla. genes* | *Scaffold* | *Orthogroup* | *D. mel. gene id* | *D. mel. gene name* | *logFC* | *adj.P.Val* | *Stage* |
|---|---|---|---|---|---|---|---|
| *jg14802* | WW_tarseq_540_arrow | OG0000503 | NP_651812.1, NP_001263113.1, NP_001189282.1 | epidermal stripes and patches, isoform B [Drosophila melanogaster] | 3.04 | 0.04 | premel |
| *jg15101* | WW_tarseq_540_arrow | OG0000975 | NP_001260005.1 | uncharacterized protein Dmel_CG43707, isoform E [Drosophila melanogaster] | -3.88 | 0 | mel |
| *jg1308* | WW_tarseq_419_arrow | OG0001450 | NP_524344.1 | yellow-e | 10.31 | 0.01 | premel |
| *jg1310* | WW_tarseq_419_arrow | OG0001450 | NP_524344.1 | yellow-e | 3.86 | 0.02 | premel |
| *jg2035* | WW_tarseq_531_arrow | OG0007293 | NP_732407.1 | cryptochrome [Drosophila melanogaster] | 2.95 | 0.05 | premel |
| *jg15103* | WW_tarseq_540_arrow | OG0007341 | NP_609535.1 | WD repeat domain 81 [Drosophila melanogaster] | 1.64 | 0.02 | 72h |
| *jg15168* | WW_tarseq_540_arrow | OG0007353 | NP_572341.1 | uncharacterized protein Dmel_CG3184 [Drosophila melanogaster] | 2.21 | 0.03 | 5days |
| *jg1153* | WW_tarseq_419_arrow | | | | -6.95 | 0 | mel |
| *jg14032* | WW_tarseq_487_arrow | | | | -1.7 | 0.04 | premel |
| *jg2034* | WW_tarseq_531_arrow | | | | 2.49 | 0.04 | premel |
| *jg8680* | WW_tarseq_472_arrow | | | | 3.42 | 0 | 5days |
| *jg9028* | WW_tarseq_472_arrow | | | | 3.24 | 0.04 | mel |

Table S4: Details of the number of eggs injected with each sgRNA and those which produced adult moths.

| sgRNAs | Number_eggs_injected | Number_larvae | Percentage_hatched | Number_pupae | Number_adults | Hatched_to_adult percentage | Percentage of emerged adults with phenotype |
|---|---|---|---|---|---|---|---|
| Val1A | 109 | 29 | 26.61 | 3 | 2 | 6.9 | 100% inc. pupa |
| Val2A | 58 | 21 | 36.21 | 0 | 0 | 0.0 | |
| Val2B | 112 | 8 | 7.14 | 1 | 1 | 12.5 | 100% (mosaic female) |
| Val3A | 216 | 11 | 5.09 | 2 | 2 | 18.2 | 50% |
| Val3B | 309 | 20 | 6.47 | 2 | 1 | 5.0 | 100% inc. pupa |
| Val1A + Val2A | 65 | 4 | 6.15 | 0 | 0 | 0.0 | |
| Val1A + Val2B | 21 | 6 | 28.57 | 0 | 0 | 0.0 | |
| Val1A + Val3A | 61 | 5 | 8.20 | 0 | 0 | 0.0 | |
| Val2A + Val2B | 55 | 9 | 16.36 | 0 | 0 | 0.0 | |
| Val2B + Val3A | 49 | 6 | 12.24 | 0 | 0 | 0.0 | |
| Val3A + Val2B | 53 | 17 | 32.08 | 0 | 0 | 0.0 | |
| Val3A + Val3B | 115 | 6 | 5.22 | 0 | 0 | 0.0 | |

Table S5: Sample list of all lab cross individuals used in linkage mapping.

| ID | Sex | Genotype | Family number | Generation | Total sequenced reads | YY mapped reads | YY % mapped reads | YY duplication | YY reads retained after deduplication | WW mapped reads | WW % mapped | WW duplication | WW reads retained after deduplication |
|---|---|---|---|---|---|---|---|---|---|---|---|---|---|
| FI17_3_34_1 | m | WY | 29 | Parent | 2955970 | 2715026 | 91.849 | 0.855 | 393347 | 2748494 | 92.98 | 0.80 | 544140 |
| FI17_3_83_11 | f | YY | 29 | Parent | 3197702 | 3136607 | 98.089 | 0.841 | 499345 | 3147370 | 98.43 | 0.76 | 823864 |
| FI18_1_29_1 | m | YY | 29 | F1 | 1999390 | 1946873 | 97.373 | 0.824 | 342770 | 1971086 | 98.58 | 0.83 | 342138 |
| FI18_1_29_101 | m | YY | 29 | F1 | 2361064 | 2248858 | 95.248 | 0.820 | 404408 | 2277871 | 96.48 | 0.82 | 404261 |
| FI18_1_29_11 | m | YY | 29 | F1 | 2068918 | 2015804 | 97.433 | 0.827 | 348902 | 2042671 | 98.73 | 0.83 | 348176 |
| FI18_1_29_19 | m | WY | 29 | F1 | 1146544 | 1115347 | 97.279 | 0.744 | 285992 | 1130950 | 98.64 | 0.75 | 285547 |
| FI18_1_29_2 | m | YY | 29 | F1 | 3529048 | 3407536 | 96.557 | 0.827 | 590114 | 3454443 | 97.89 | 0.83 | 591096 |
| FI18_1_29_23 | m | YY | 29 | F1 | 1283616 | 1242794 | 96.820 | 0.740 | 323494 | 1259702 | 98.14 | 0.74 | 323404 |
| FI18_1_29_27 | m | YY | 29 | F1 | 2399278 | 2331282 | 97.166 | 0.788 | 493892 | 2364010 | 98.53 | 0.79 | 494821 |
| FI18_1_29_32 | m | YY | 29 | F1 | 1816056 | 1767863 | 97.346 | 0.816 | 325860 | 1789547 | 98.54 | 0.82 | 324933 |
| FI18_1_29_33 | m | YY | 29 | F1 | 2164252 | 2100526 | 97.056 | 0.768 | 487796 | 2130653 | 98.45 | 0.77 | 488153 |
| FI18_1_29_38 | m | WY | 29 | F1 | 3748912 | 3647923 | 97.306 | 0.870 | 473781 | 3697017 | 98.62 | 0.87 | 471901 |
| FI18_1_29_39 | m | YY | 29 | F1 | 1695450 | 1237858 | 73.011 | 0.740 | 321648 | 1254382 | 73.99 | 0.74 | 321608 |
| FI18_1_29_4 | m | WY | 29 | F1 | 1659596 | 1617162 | 97.443 | 0.810 | 307254 | 1637651 | 98.68 | 0.81 | 306746 |
| FI18_1_29_41 | m | WY | 29 | F1 | 1067638 | 1035473 | 96.987 | 0.703 | 307780 | 1050028 | 98.35 | 0.71 | 307601 |
| FI18_1_29_43 | m | WY | 29 | F1 | 1180652 | 1130239 | 95.730 | 0.756 | 276252 | 1145620 | 97.03 | 0.76 | 275715 |
| FI18_1_29_44 | m | WY | 29 | F1 | 2891056 | 2818722 | 97.498 | 0.820 | 506972 | 2854834 | 98.75 | 0.82 | 507386 |
| FI18_1_29_46 | m | YY | 29 | F1 | 3147978 | 3067342 | 97.438 | 0.820 | 551441 | 3107076 | 98.70 | 0.82 | 552171 |
| FI18_1_29_49 | m | WY | 29 | F1 | 794124 | 770168 | 96.983 | 0.686 | 241694 | 780562 | 98.29 | 0.69 | 241493 |
| FI18_1_29_58 | m | YY | 29 | F1 | 2893010 | 1473576 | 50.936 | 0.706 | 432618 | 1496051 | 51.71 | 0.71 | 433794 |
| FI18_1_29_6 | m | YY | 29 | F1 | 3159060 | 3034452 | 96.056 | 0.814 | 564750 | 3077094 | 97.41 | 0.82 | 565680 |
| FI18_1_29_60 | m | WY | 29 | F1 | 1415616 | 1369725 | 96.758 | 0.762 | 326207 | 1388092 | 98.06 | 0.77 | 326034 |
| FI18_1_29_65 | m | YY | 29 | F1 | 3140294 | 3051117 | 97.160 | 0.841 | 484739 | 3093059 | 98.50 | 0.84 | 484175 |
| FI18_1_29_68 | m | YY | 29 | F1 | 2696202 | 2626553 | 97.417 | 0.855 | 381758 | 2657543 | 98.57 | 0.86 | 380774 |
| FI18_1_29_74 | m | YY | 29 | F1 | 1449948 | 1413194 | 97.465 | 0.777 | 314744 | 1431318 | 98.72 | 0.78 | 313807 |
| FI18_1_29_76 | m | YY | 29 | F1 | 2531304 | 2418112 | 95.528 | 0.853 | 355571 | 2447038 | 96.67 | 0.85 | 354981 |
| FI18_1_29_78 | m | YY | 29 | F1 | 1678066 | 1625019 | 96.839 | 0.779 | 359452 | 1647238 | 98.16 | 0.78 | 359548 |
| FI18_1_29_82 | m | YY | 29 | F1 | 3198008 | 3097633 | 96.861 | 0.858 | 439408 | 3136758 | 98.08 | 0.86 | 438710 |
| FI18_1_29_83 | m | YY | 29 | F1 | 3006038 | 2866236 | 95.349 | 0.830 | 486751 | 2907742 | 96.73 | 0.83 | 485152 |
| FI18_1_29_84 | m | WY | 29 | F1 | 1520480 | 1481081 | 97.409 | 0.789 | 312609 | 1499436 | 98.62 | 0.79 | 312154 |
| FI18_1_29_88 | m | YY | 29 | F1 | 2657610 | 2587124 | 97.348 | 0.822 | 460265 | 2620810 | 98.62 | 0.82 | 460215 |
| FI18_1_29_9 | m | WY | 29 | F1 | 1554302 | 1504346 | 96.786 | 0.772 | 343716 | 1525473 | 98.15 | 0.77 | 343466 |

| | | | | | | | | | | | | | |
|---|---|---|---|---|---|---|---|---|---|---|---|---|---|
| FI18_1_29_93 | m | WY | 29 | F1 | 1737742 | 1691286 | 97.327 | 0.810 | 321434 | 1712318 | 98.54 | 0.81 | 320585 |
| FI18_1_29_95 | m | WY | 29 | F1 | 2188534 | 2058702 | 94.068 | 0.815 | 381273 | 2085456 | 95.29 | 0.82 | 380670 |
| FI18_1_29_98 | m | YY | 29 | F1 | 2821014 | 1951884 | 69.191 | 0.755 | 478442 | 1980282 | 70.20 | 0.76 | 479244 |
| FI17_3_71_3 | m | WY | 30 | Parent | 4317516 | 4162243 | 96.404 | 0.853 | 613865 | 4224077 | 97.84 | 0.80 | 830911 |
| FI17_3_81_4 | f | YY | 30 | Parent | 3107120 | 3026320 | 97.400 | 0.857 | 433556 | 3040627 | 97.86 | 0.75 | 541186 |
| FI18_1_30_106 | m | WY | 30 | F1 | 2715010 | 2053347 | 75.629 | 0.790 | 431541 | 2083411 | 76.74 | 0.79 | 431795 |
| FI18_1_30_11 | m | WY | 30 | F1 | 2869392 | 2778003 | 96.815 | 0.848 | 423555 | 2815377 | 98.12 | 0.85 | 422981 |
| FI18_1_30_12 | m | WY | 30 | F1 | 2973814 | 2844480 | 95.651 | 0.834 | 473217 | 2886012 | 97.05 | 0.84 | 471277 |
| FI18_1_30_14 | m | WY | 30 | F1 | 4478234 | 2564695 | 57.270 | 0.832 | 431954 | 2601818 | 58.10 | 0.83 | 430023 |
| FI18_1_30_18 | m | WY | 30 | F1 | 1599220 | 1555974 | 97.296 | 0.818 | 283223 | 1577278 | 98.63 | 0.82 | 281921 |
| FI18_1_30_2 | m | YY | 30 | F1 | 3022868 | 2785048 | 92.133 | 0.823 | 492599 | 2823454 | 93.40 | 0.83 | 492254 |
| FI18_1_30_24 | m | WY | 30 | F1 | 1825228 | 1516006 | 83.058 | 0.794 | 311592 | 1538155 | 84.27 | 0.80 | 310094 |
| FI18_1_30_3 | m | YY | 30 | F1 | 1734848 | 1022616 | 58.946 | 0.809 | 195239 | 1034509 | 59.63 | 0.81 | 193181 |
| FI18_1_30_30 | m | WY | 30 | F1 | 2787488 | 2698920 | 96.823 | 0.801 | 536756 | 2740534 | 98.32 | 0.80 | 535060 |
| FI18_1_30_32 | m | YY | 30 | F1 | 2379786 | 2315421 | 97.295 | 0.827 | 400142 | 2349891 | 98.74 | 0.83 | 398925 |
| FI18_1_30_36 | m | WY | 30 | F1 | 1527360 | 1339985 | 87.732 | 0.774 | 303226 | 1359367 | 89.00 | 0.78 | 302162 |
| FI18_1_30_4 | m | WY | 30 | F1 | 2383670 | 2321238 | 97.381 | 0.836 | 381226 | 2352819 | 98.71 | 0.84 | 379064 |
| FI18_1_30_40 | m | WY | 30 | F1 | 2346222 | 2284549 | 97.371 | 0.843 | 359175 | 2315827 | 98.70 | 0.85 | 356863 |
| FI18_1_30_42 | m | YY | 30 | F1 | 1070592 | 1038205 | 96.975 | 0.768 | 241165 | 1053339 | 98.39 | 0.77 | 240238 |
| FI18_1_30_43 | m | YY | 30 | F1 | 1972550 | 1918571 | 97.263 | 0.821 | 343546 | 1946119 | 98.66 | 0.82 | 342191 |
| FI18_1_30_5 | m | WY | 30 | F1 | 1426882 | 1367090 | 95.810 | 0.788 | 290415 | 1387392 | 97.23 | 0.79 | 289031 |
| FI18_1_30_52 | m | WY | 30 | F1 | 5053696 | 4514518 | 89.331 | 0.892 | 487534 | 4574384 | 90.52 | 0.89 | 481237 |
| FI18_1_30_53 | m | YY | 30 | F1 | 2230112 | 2171645 | 97.378 | 0.818 | 396180 | 2196873 | 98.51 | 0.82 | 397275 |
| FI18_1_30_54 | m | YY | 30 | F1 | 3919162 | 3799033 | 96.935 | 0.883 | 443179 | 3851217 | 98.27 | 0.89 | 436759 |
| FI18_1_30_55 | m | WY | 30 | F1 | 2322312 | 2192761 | 94.421 | 0.818 | 399266 | 2223184 | 95.73 | 0.82 | 398852 |
| FI18_1_30_56 | m | WY | 30 | F1 | 1114538 | 1082869 | 97.159 | 0.733 | 288835 | 1098673 | 98.58 | 0.74 | 288058 |
| FI18_1_30_57 | m | WY | 30 | F1 | 3931316 | 3777836 | 96.096 | 0.868 | 500346 | 3827592 | 97.36 | 0.87 | 496429 |
| FI18_1_30_62 | m | WY | 30 | F1 | 2868180 | 2791424 | 97.324 | 0.835 | 459977 | 2830970 | 98.70 | 0.84 | 456986 |
| FI18_1_30_63 | m | YY | 30 | F1 | 1461576 | 1423021 | 97.362 | 0.790 | 299500 | 1441196 | 98.61 | 0.79 | 299667 |
| FI18_1_30_65 | m | WY | 30 | F1 | 1866314 | 1812580 | 97.121 | 0.827 | 314087 | 1838963 | 98.53 | 0.83 | 312070 |
| FI18_1_30_66 | m | WY | 30 | F1 | 3043110 | 2936318 | 96.491 | 0.896 | 305921 | 2970116 | 97.60 | 0.90 | 301052 |
| FI18_1_30_75 | m | WY | 30 | F1 | 2295070 | 2216385 | 96.572 | 0.813 | 414891 | 2249790 | 98.03 | 0.82 | 413818 |
| FI18_1_30_8 | m | WY | 30 | F1 | 3093078 | 2659531 | 85.983 | 0.789 | 562165 | 2701905 | 87.35 | 0.79 | 561879 |
| FI18_1_30_80 | m | WY | 30 | F1 | 2270474 | 2167775 | 95.477 | 0.785 | 465533 | 2201415 | 96.96 | 0.79 | 464124 |
| FI18_1_30_84 | m | WY | 30 | F1 | 3155708 | 3061936 | 97.028 | 0.832 | 515203 | 3109723 | 98.54 | 0.84 | 512912 |
| FI18_1_30_88 | m | WY | 30 | F1 | 2060846 | 1877502 | 91.103 | 0.831 | 318137 | 1904332 | 92.41 | 0.83 | 316669 |
| FI18_1_30_89 | m | WY | 30 | F1 | 4167516 | 3948027 | 94.733 | 0.859 | 557287 | 4006846 | 96.14 | 0.86 | 553226 |
| FI18_1_30_9 | m | WY | 30 | F1 | 145926 | 139007 | 95.259 | 0.468 | 73949 | 141042 | 96.65 | 0.48 | 73822 |

| | | | | | | | | | | | | | |
|---|---|---|---|---|---|---|---|---|---|---|---|---|---|
| FI18_1_30_90 | m | YY | 30 | F1 | 1286248 | 1252107 | 97.346 | 0.767 | 291522 | 1270035 | 98.74 | 0.77 | 291225 |
| FI18_1_30_91 | m | WY | 30 | F1 | 2326852 | 2245625 | 96.509 | 0.816 | 412438 | 2279310 | 97.96 | 0.82 | 410759 |
| FI18_1_30_92 | m | YY | 30 | F1 | 1830454 | 1770263 | 96.712 | 0.769 | 408900 | 1796330 | 98.14 | 0.77 | 408877 |
| FI18_1_30_94 | m | YY | 30 | F1 | 3836556 | 3698968 | 96.414 | 0.871 | 478601 | 3743031 | 97.56 | 0.87 | 479708 |
| FI18_1_30_95 | m | WY | 30 | F1 | 4085302 | 3956235 | 96.841 | 0.873 | 501017 | 4013996 | 98.25 | 0.88 | 495944 |
| FI17_3_97_7 | m | WY | 32 | Parent | 5704750 | 3351852 | 58.755 | 0.843 | 525582 | 3400691 | 59.61 | 0.80 | 503529 |
| FI17_3_56_2 | f | YY | 32 | Parent | 12734726 | 2216899 | 17.408 | 0.856 | 318198 | 2224846 | 17.47 | 0.80 | 445434 |
| FI18_1_32_1 | m | WY | 32 | F1 | 2713466 | 2628102 | 96.854 | 0.853 | 386373 | 2661104 | 98.07 | 0.86 | 384355 |
| FI18_1_32_10 | m | WY | 32 | F1 | 3211340 | 3131817 | 97.524 | 0.859 | 440030 | 3169547 | 98.70 | 0.86 | 437076 |
| FI18_1_32_100 | m | WY | 32 | F1 | 2651298 | 2586070 | 97.540 | 0.849 | 390910 | 2619419 | 98.80 | 0.85 | 387892 |
| FI18_1_32_11 | m | WY | 32 | F1 | 2015790 | 1953474 | 96.909 | 0.804 | 382149 | 1979883 | 98.22 | 0.81 | 380754 |
| FI18_1_32_12 | m | YY | 32 | F1 | 2420446 | 2339031 | 96.636 | 0.807 | 452120 | 2371340 | 97.97 | 0.81 | 450573 |
| FI18_1_32_17 | m | WY | 32 | F1 | 2503044 | 2419974 | 96.681 | 0.852 | 358779 | 2450754 | 97.91 | 0.85 | 356650 |
| FI18_1_32_18 | m | YY | 32 | F1 | 1672500 | 1628328 | 97.359 | 0.817 | 297256 | 1648404 | 98.56 | 0.82 | 295997 |
| FI18_1_32_23 | m | YY | 32 | F1 | 2664986 | 2484766 | 93.237 | 0.848 | 377868 | 2516373 | 94.42 | 0.85 | 375861 |
| FI18_1_32_27 | m | YY | 32 | F1 | 2594498 | 2525197 | 97.329 | 0.823 | 447059 | 2559795 | 98.66 | 0.83 | 445908 |
| FI18_1_32_30 | m | YY | 32 | F1 | 1720144 | 1675773 | 97.421 | 0.814 | 311376 | 1697540 | 98.69 | 0.82 | 310141 |
| FI18_1_32_36 | m | WY | 32 | F1 | 2220868 | 2156152 | 97.086 | 0.781 | 473166 | 2186450 | 98.45 | 0.78 | 473939 |
| FI18_1_32_38 | m | WY | 32 | F1 | 2945502 | 2866956 | 97.333 | 0.853 | 421535 | 2906212 | 98.67 | 0.86 | 418895 |
| FI18_1_32_40 | m | WY | 32 | F1 | 1945642 | 1737773 | 89.316 | 0.798 | 350668 | 1761860 | 90.55 | 0.80 | 349957 |
| FI18_1_32_44 | m | YY | 32 | F1 | 2912328 | 2813114 | 96.593 | 0.856 | 406172 | 2849472 | 97.84 | 0.86 | 402462 |
| FI18_1_32_47 | m | WY | 32 | F1 | 1805192 | 1568781 | 86.904 | 0.771 | 360027 | 1591403 | 88.16 | 0.77 | 360198 |
| FI18_1_32_50_1 | m | WY | 32 | F1 | 2343302 | 2267978 | 96.786 | 0.838 | 367425 | 2296497 | 98.00 | 0.84 | 365773 |
| FI18_1_32_51 | m | WY | 32 | F1 | 2274740 | 2089964 | 91.877 | 0.833 | 348148 | 2115979 | 93.02 | 0.84 | 346551 |
| FI18_1_32_53 | m | YY | 32 | F1 | 1364862 | 1317458 | 96.527 | 0.774 | 297759 | 1334444 | 97.77 | 0.78 | 296711 |
| FI18_1_32_56 | m | YY | 32 | F1 | 2223158 | 2018476 | 90.793 | 0.837 | 328854 | 2044373 | 91.96 | 0.84 | 326572 |
| FI18_1_32_57 | m | YY | 32 | F1 | 2584886 | 2491294 | 96.379 | 0.833 | 416906 | 2523897 | 97.64 | 0.84 | 415749 |
| FI18_1_32_58 | m | YY | 32 | F1 | 2042488 | 1987407 | 97.303 | 0.804 | 389592 | 2015607 | 98.68 | 0.81 | 388087 |
| FI18_1_32_66 | m | WY | 32 | F1 | 2861330 | 2734683 | 95.574 | 0.817 | 500515 | 2772097 | 96.88 | 0.82 | 499501 |
| FI18_1_32_67 | m | YY | 32 | F1 | 2683192 | 2573925 | 95.928 | 0.841 | 408704 | 2607463 | 97.18 | 0.84 | 406990 |
| FI18_1_32_68 | m | YY | 32 | F1 | 2192898 | 2138659 | 97.527 | 0.830 | 363067 | 2165772 | 98.76 | 0.83 | 361189 |
| FI18_1_32_70 | m | YY | 32 | F1 | 2519412 | 2446488 | 97.106 | 0.793 | 507448 | 2482049 | 98.52 | 0.80 | 507378 |
| FI18_1_32_71 | m | YY | 32 | F1 | 3308020 | 3171685 | 95.879 | 0.851 | 472947 | 3212971 | 97.13 | 0.85 | 470874 |
| FI18_1_32_78 | m | WY | 32 | F1 | 2112880 | 2062840 | 97.632 | 0.839 | 332683 | 2088674 | 98.85 | 0.84 | 330912 |
| FI18_1_32_79 | m | YY | 32 | F1 | 2518434 | 1915805 | 76.071 | 0.812 | 360674 | 1942127 | 77.12 | 0.81 | 359605 |
| FI18_1_32_80 | m | WY | 32 | F1 | 1477900 | 1224353 | 82.844 | 0.771 | 280331 | 1241392 | 84.00 | 0.77 | 279709 |
| FI18_1_32_83 | m | WY | 32 | F1 | 1117270 | 1078079 | 96.492 | 0.731 | 289881 | 1093619 | 97.88 | 0.73 | 290086 |
| FI18_1_32_88 | m | YY | 32 | F1 | 2058940 | 2004230 | 97.343 | 0.826 | 347869 | 2029105 | 98.55 | 0.83 | 346436 |

| | | | | | | | | | | | | | |
|---|---|---|---|---|---|---|---|---|---|---|---|---|---|
| FI18_1_32_89 | m | WY | 32 | F1 | 2449154 | 2384743 | 97.370 | 0.824 | 420528 | 2414835 | 98.60 | 0.83 | 418755 |
| FI18_1_32_9 | m | YY | 32 | F1 | 2106886 | 2039571 | 96.805 | 0.835 | 336647 | 2064595 | 97.99 | 0.84 | 334520 |
| FI17_3_72_7 | m | WY | 35 | Parent | 6985806 | 6733807 | 96.393 | 0.918 | 555217 | 6820593 | 97.64 | 0.89 | 781200 |
| FI17_3_49_2 | f | YY | 35 | Parent | 4170902 | 3996659 | 95.822 | 0.881 | 474906 | 4008431 | 96.10 | 0.84 | 653585 |
| FI18_1_35_105 | m | YY | 35 | F1 | 4022634 | 3834980 | 95.335 | 0.843 | 602628 | 3889267 | 96.68 | 0.85 | 600085 |
| FI18_1_35_106 | m | YY | 35 | F1 | 3614566 | 3030045 | 83.829 | 0.834 | 502926 | 3071448 | 84.97 | 0.84 | 501637 |
| FI18_1_35_107 | m | YY | 35 | F1 | 3346384 | 3254695 | 97.260 | 0.861 | 451300 | 3297645 | 98.54 | 0.86 | 448886 |
| FI18_1_35_108 | m | WY | 35 | F1 | 2368662 | 2285847 | 96.504 | 0.827 | 394503 | 2316049 | 97.78 | 0.83 | 393746 |
| FI18_1_35_109 | m | YY | 35 | F1 | 4079696 | 3966821 | 97.233 | 0.853 | 581428 | 4020088 | 98.54 | 0.86 | 578209 |
| FI18_1_35_11 | m | WY | 35 | F1 | 2111094 | 2000420 | 94.758 | 0.826 | 414635 | 2026564 | 96.00 | 0.83 | 352689 |
| FI18_1_35_110 | m | WY | 35 | F1 | 2458458 | 2389520 | 97.196 | 0.826 | 386197 | 2421034 | 98.48 | 0.83 | 414091 |
| FI18_1_35_111 | m | WY | 35 | F1 | 2303368 | 2222441 | 96.487 | 0.823 | 354631 | 2252832 | 97.81 | 0.83 | 384555 |
| FI18_1_35_112 | m | WY | 35 | F1 | 3711582 | 3584681 | 96.581 | 0.866 | 479803 | 3630766 | 97.82 | 0.87 | 476672 |
| FI18_1_35_113 | m | WY | 35 | F1 | 3159130 | 3069909 | 97.176 | 0.829 | 524186 | 3113018 | 98.54 | 0.83 | 522785 |
| FI18_1_35_114 | m | WY | 35 | F1 | 2286442 | 2219116 | 97.055 | 0.796 | 451671 | 2251683 | 98.48 | 0.80 | 451709 |
| FI18_1_35_115 | m | YY | 35 | F1 | 4042534 | 3556216 | 87.970 | 0.908 | 328073 | 3596944 | 88.98 | 0.91 | 323326 |
| FI18_1_35_117 | m | YY | 35 | F1 | 5095106 | 4910918 | 96.385 | 0.877 | 605879 | 4976003 | 97.66 | 0.88 | 607205 |
| FI18_1_35_118 | m | WY | 35 | F1 | 2803240 | 2713436 | 96.796 | 0.837 | 441079 | 2749086 | 98.07 | 0.84 | 439911 |
| FI18_1_35_12 | m | YY | 35 | F1 | 5122120 | 4318359 | 84.308 | 0.892 | 465882 | 4367086 | 85.26 | 0.89 | 460939 |
| FI18_1_35_124 | m | YY | 35 | F1 | 4084550 | 3981723 | 97.483 | 0.893 | 425219 | 4031405 | 98.70 | 0.90 | 419245 |
| FI18_1_35_129 | m | WY | 35 | F1 | 1868866 | 1812576 | 96.988 | 0.823 | 320517 | 1837315 | 98.31 | 0.83 | 318853 |
| FI18_1_35_130 | m | WY | 35 | F1 | 2278784 | 2220546 | 97.444 | 0.851 | 330748 | 2249967 | 98.74 | 0.85 | 328178 |
| FI18_1_35_131 | m | YY | 35 | F1 | 2508666 | 2438390 | 97.199 | 0.839 | 391789 | 2469826 | 98.45 | 0.84 | 390368 |
| FI18_1_35_14 | m | YY | 35 | F1 | 5116298 | 4944259 | 96.637 | 0.859 | 697050 | 5013442 | 97.99 | 0.86 | 694775 |
| FI18_1_35_15 | m | YY | 35 | F1 | 3019894 | 2890565 | 95.717 | 0.785 | 621455 | 2933036 | 97.12 | 0.79 | 622407 |
| FI18_1_35_16 | m | WY | 35 | F1 | 2456980 | 2387996 | 97.192 | 0.833 | 397605 | 2421394 | 98.55 | 0.84 | 395404 |
| FI18_1_35_18 | m | YY | 35 | F1 | 3912406 | 3805044 | 97.256 | 0.885 | 436710 | 3854079 | 98.51 | 0.89 | 434097 |
| FI18_1_35_201 | m | YY | 35 | F1 | 3533726 | 3429725 | 97.057 | 0.862 | 472487 | 3475867 | 98.36 | 0.86 | 470227 |
| FI18_1_35_202 | m | YY | 35 | F1 | 1548800 | 1497770 | 96.705 | 0.789 | 315985 | 1519757 | 98.12 | 0.79 | 315829 |
| FI18_1_35_203 | m | WY | 35 | F1 | 2529636 | 2361197 | 93.341 | 0.815 | 435746 | 2394939 | 94.68 | 0.82 | 434933 |
| FI18_1_35_204 | m | WY | 35 | F1 | 1843578 | 1688393 | 91.582 | 0.763 | 400723 | 1713748 | 92.96 | 0.77 | 401016 |
| FI18_1_35_205 | m | WY | 35 | F1 | 2985074 | 2906901 | 97.381 | 0.864 | 396332 | 2944441 | 98.64 | 0.87 | 393787 |
| FI18_1_35_23 | m | WY | 35 | F1 | 2660236 | 2517194 | 94.623 | 0.845 | 390549 | 2550049 | 95.86 | 0.85 | 388521 |
| FI18_1_35_26 | m | WY | 35 | F1 | 3518410 | 3419253 | 97.182 | 0.848 | 521155 | 3465568 | 98.50 | 0.85 | 519575 |
| FI18_1_35_27 | m | YY | 35 | F1 | 4180740 | 4018770 | 96.126 | 0.859 | 566600 | 4073081 | 97.42 | 0.86 | 564290 |
| FI18_1_35_30 | m | YY | 35 | F1 | 3154926 | 3052328 | 96.748 | 0.883 | 356336 | 3091011 | 97.97 | 0.89 | 351995 |
| FI18_1_35_34 | m | YY | 35 | F1 | 3944846 | 3793771 | 96.170 | 0.857 | 540650 | 3844661 | 97.46 | 0.86 | 539733 |
| FI18_1_35_38 | m | YY | 35 | F1 | 2862154 | 2762897 | 96.532 | 0.861 | 383162 | 2799925 | 97.83 | 0.86 | 380537 |

| | | | | | | | | | | | | | |
|---|---|---|---|---|---|---|---|---|---|---|---|---|---|
| FI18_1_35_4 | m | WY | 35 | F1 | 1577512 | 1536243 | 97.384 | 0.808 | 295506 | 1555941 | 98.63 | 0.87 | 438536 |
| FI18_1_35_41 | m | YY | 35 | F1 | 3559782 | 3367393 | 94.595 | 0.869 | 441184 | 3410928 | 95.82 | 0.81 | 293362 |
| FI18_1_35_42 | m | YY | 35 | F1 | 2472706 | 2299419 | 92.992 | 0.846 | 353007 | 2327801 | 94.14 | 0.85 | 351126 |
| FI18_1_35_44 | m | WY | 35 | F1 | 2710278 | 2619093 | 96.636 | 0.842 | 414429 | 2655136 | 97.97 | 0.84 | 411613 |
| FI18_1_35_46 | m | YY | 35 | F1 | 4262616 | 4140732 | 97.141 | 0.862 | 570989 | 4198044 | 98.49 | 0.86 | 570002 |
| FI18_1_35_48 | m | WY | 35 | F1 | 3216496 | 2916095 | 90.661 | 0.856 | 421370 | 2953800 | 91.83 | 0.86 | 418047 |
| FI18_1_35_50 | m | WY | 35 | F1 | 2489832 | 2423228 | 97.325 | 0.848 | 368727 | 2455548 | 98.62 | 0.85 | 366764 |
| FI18_1_35_55 | m | WY | 35 | F1 | 2359414 | 2275714 | 96.453 | 0.820 | 408809 | 2306363 | 97.75 | 0.82 | 407863 |
| FI18_1_35_63 | m | WY | 35 | F1 | 2052154 | 1974678 | 96.225 | 0.825 | 344993 | 2000431 | 97.48 | 0.83 | 343929 |
| FI18_1_35_64 | m | WY | 35 | F1 | 1527192 | 1485772 | 97.288 | 0.782 | 323976 | 1505656 | 98.59 | 0.79 | 323437 |
| FI18_1_35_7 | m | YY | 35 | F1 | 5425568 | 5283782 | 97.387 | 0.888 | 593717 | 5352263 | 98.65 | 0.89 | 588934 |
| FI18_1_35_71 | m | WY | 35 | F1 | 2400100 | 2329217 | 97.047 | 0.819 | 421606 | 2360482 | 98.35 | 0.82 | 420247 |
| FI18_1_35_75 | m | WY | 35 | F1 | 964512 | 751542 | 77.919 | 0.778 | 166550 | 758586 | 78.65 | 0.78 | 165028 |
| FI18_1_35_77 | m | YY | 35 | F1 | 3730104 | 3620632 | 97.065 | 0.865 | 488724 | 3666177 | 98.29 | 0.87 | 488414 |
| FI18_1_35_8 | m | YY | 35 | F1 | 3452032 | 3358858 | 97.301 | 0.846 | 518108 | 3403069 | 98.58 | 0.85 | 515878 |
| FI18_1_35_80 | m | WY | 35 | F1 | 2593944 | 2496331 | 96.237 | 0.818 | 454804 | 2533709 | 97.68 | 0.82 | 453525 |
| FI18_1_35_87 | m | YY | 35 | F1 | 1765316 | 1719243 | 97.390 | 0.820 | 308853 | 1741749 | 98.66 | 0.82 | 307339 |
| FI18_1_35_88 | m | YY | 35 | F1 | 4767910 | 4573602 | 95.925 | 0.866 | 611810 | 4636650 | 97.25 | 0.87 | 607947 |
| FI18_1_35_90 | m | YY | 35 | F1 | 3354264 | 3252455 | 96.965 | 0.878 | 395524 | 3293166 | 98.18 | 0.88 | 392171 |
| FI18_1_35_95 | m | YY | 35 | F1 | 4430800 | 4302387 | 97.102 | 0.842 | 678160 | 4362196 | 98.45 | 0.84 | 678885 |
| FI18_1_35_96 | m | WY | 35 | F1 | 2715398 | 2640153 | 97.229 | 0.835 | 436057 | 2676128 | 98.55 | 0.84 | 435103 |
| FI18_1_35_97 | m | WY | 35 | F1 | 4886316 | 4653904 | 95.244 | 0.899 | 471477 | 4712210 | 96.44 | 0.90 | 467161 |
| FI18_1_29_14_1 | excluded due to low mapping | | 29 | F1 | 5754900 | 274244 | 4.765 | | | 278459 | 4.84 | | |
| FI18_1_29_20_1 | excluded due to low mapping | | 29 | F1 | 691934 | 153766 | 22.223 | | | 155963 | 22.54 | | |
| FI18_1_29_3_1 | excluded due to low mapping | | 29 | F1 | 6673428 | 349921 | 5.243 | | | 352906 | 5.29 | | |
| FI18_1_29_59_1 | excluded due to low mapping | | 29 | F1 | 2940246 | 206153 | 7.011 | | | 208804 | 7.10 | | |
| FI18_1_29_97_1 | excluded due to low mapping | | 29 | F1 | 7207076 | 1488963 | 20.660 | | | 1508887 | 20.94 | | |
| FI18_1_30_96_1 | excluded due to low mapping | | 30 | F1 | 1115724 | | | | | | | | |
| FI18_1_32_50_2_1 | excluded due to low mapping | | 32 | F1 | 9689116 | 368989 | 3.808 | | | 373251 | 3.85 | | |
| FI18_1_32_74_1 | excluded due to low mapping | | 32 | F1 | 6430210 | 1215692 | 18.906 | | | 1229932 | 19.13 | | |
| FI18_1_32_94_1 | excluded due to low mapping | | 32 | F1 | 6933950 | 5839 | 0.084 | | | 5963 | 0.09 | | |
| FI18_1_35_53_1 | excluded due to low mapping | | 35 | F1 | 11105008 | 10313 | 0.093 | | | 10423 | 0.09 | | |
| FI18_1_35_91_1 | excluded due to low mapping | | 35 | F1 | 9853700 | 62965 | 0.639 | | | 63712 | 0.65 | | |
| FI18_1_35_94_1 | excluded due to low mapping | | 35 | F1 | 6052848 | 1562498 | 25.814 | | | 1582649 | 26.15 | | |

Table S6: Sample list of all wild samples used.

| ID | Locality | Latitude | Longitude | Region | Collection year | Sex | Colour | Used in GWAS | Total sequenced reads | YY Mapped reads | YY % mapped reads | YY duplication | YY Reads retained after deduplication | YY average read depth | WW mapped read | WW % mapped | WW duplication | WW Reads retained after deduplication | WW average read depth |
|---|---|---|---|---|---|---|---|---|---|---|---|---|---|---|---|---|---|---|---|
| CAM015132 | Huosiaisnotko, Laukaa | 62.38634 | 25.81796 | Central Finland | 2018 | Male | Y | x | 99337203 | 95728163 | 96.37 | 0.13 | 83260117 | 17.382 | 99278718 | 99.02 | 0.14 | 98307916 | 17.6129 |
| **CAM015133** | Huosiaisnotko, Laukaa | 62.38634 | 25.81796 | Central Finland | 2018 | Male | Y | x | 91549356 | 88141528 | 96.28 | 0.14 | 75592171 | 15.8571 | 91510105 | 98.94 | 0.15 | 90540109 | 16.0169 |
| **CAM015134** | Haralanharju, Kangasala | 61.53456355 | 24.08097897 | Central Finland | 2018 | Male | Y | x | 84051672 | 81125511 | 96.52 | 0.14 | 69937864 | 14.6407 | 84005409 | 99.03 | 0.14 | 83188228 | 14.8749 |
| **CAM015135** | Heposuo, Laukaa | 62.34784 | 25.82785 | Central Finland | 2018 | Male | Y | x | 82188518 | 79203877 | 96.37 | 0.14 | 68088328 | 14.2128 | 82119542 | 99 | 0.15 | 81299902 | 14.3792 |
| **CAM015136** | Huosiaisnotko, Laukaa | 62.38634 | 25.81796 | Central Finland | 2018 | Male | Y | x | 83831177 | 80915884 | 96.52 | 0.14 | 69802421 | 14.6376 | 83766436 | 99.03 | 0.14 | 82952260 | 14.842 |
| **CAM015137** | Mäyrämäki, Jyväskylä | 62.22814 | 25.64794 | Central Finland | 2018 | Male | W | x | 90325260 | 86804526 | 96.10 | 0.14 | 74784341 | 15.8328 | 90270209 | 98.99 | 0.14 | 89355999 | 16.0069 |
| **CAM015138** | Heposuo, Laukaa | 62.34784 | 25.82785 | Central Finland | 2018 | Male | W | x | 83258916 | 80277374 | 96.42 | 0.14 | 69295411 | 14.4791 | 83204225 | 98.98 | 0.14 | 82355152 | 14.675 |
| **CAM015139** | Lautaperä, Keuruu | 62.18709 | 24.87121 | Central Finland | 2018 | Male | W | x | 91929373 | 88159990 | 95.90 | 0.14 | 75864499 | 15.7166 | 91852724 | 99.04 | 0.15 | 90968838 | 15.9305 |
| **CAM015140** | Heposuo, Laukaa | 62.34784 | 25.82785 | Central Finland | 2018 | Male | W | x | 85888608 | 82609774 | 96.18 | 0.14 | 71419687 | 14.8998 | 85825617 | 98.96 | 0.14 | 84937070 | 15.1099 |
| **CAM015141** | Huosiaisnotko, Laukaa | 62.38634 | 25.81796 | Central Finland | 2018 | Male | W | x | 76588254 | 73780022 | 96.33 | 0.14 | 63477797 | 13.3699 | 76558659 | 99.01 | 0.15 | 75803292 | 13.5154 |
| **CAM015192** | Heposuo, Laukaa | 62.34784 | 25.82785 | Central Finland | 2018 | Male | Y | x | 80404515 | 77653009 | 96.58 | 0.15 | 65662866 | 13.7736 | 80351585 | 98.85 | 0.16 | 79428391 | 14.0112 |
| **CAM015193** | Heposuo, Laukaa | 62.34784 | 25.82785 | Central Finland | 2018 | Male | Y | x | 58498076 | 56522266 | 96.62 | 0.13 | 49442117 | 10.4086 | 58462162 | 99.04 | 0.13 | 57903161 | 10.5802 |
| **CAM015194** | Heposuo, Laukaa | 62.34784 | 25.82785 | Central Finland | 2018 | Male | Y | x | 65622607 | 63282509 | 96.43 | 0.14 | 54683278 | 11.5851 | 65581436 | 98.98 | 0.14 | 64914704 | 11.7449 |
| **CAM015195** | Mäyrämäki, Jyväskylä | 62.22814 | 25.64794 | Central Finland | 2018 | Male | Y | x | 58154455 | 56272290 | 96.76 | 0.12 | 49706191 | 10.5543 | 58119500 | 99.02 | 0.12 | 57547059 | 10.6945 |
| **CAM015196** | Mäyrämäki, Jyväskylä | 62.22814 | 25.64794 | Central Finland | 2018 | Male | Y | x | 83371214 | 80690761 | 96.78 | 0.14 | 69714679 | 14.7052 | 83320131 | 99.05 | 0.14 | 82530724 | 14.8955 |
| **CAM015197** | Mäyrämäki, Jyväskylä | 62.22814 | 25.64794 | Central Finland | 2018 | Male | W | x | 68675694 | 66496196 | 96.83 | 0.13 | 58102548 | 12.2659 | 68633799 | 99.08 | 0.13 | 68004964 | 12.4399 |
| **CAM015198** | Mäyrämäki, Jyväskylä | 62.22814 | 25.64794 | Central Finland | 2018 | Male | W | x | 75453002 | 72010183 | 95.44 | 0.24 | 55086736 | 11.385 | 75405484 | 98.87 | 0.25 | 74551498 | 11.5827 |
| **CAM015199** | Mäyrämäki, Jyväskylä | 62.22814 | 25.64794 | Central Finland | 2018 | Male | W | x | 69939593 | 67389470 | 96.35 | 0.14 | 58195017 | 12.5471 | 69889281 | 99.04 | 0.14 | 69214869 | 12.7278 |
| **CAM015200** | Mäyrämäki, Jyväskylä | 62.22814 | 25.64794 | Central Finland | 2018 | Male | W | x | 55198189 | 53350748 | 96.65 | 0.12 | 47208830 | 10.2012 | 56405976 | 99.08 | 0.12 | 54666270 | 10.3219 |
| **CAM015201** | Mäyrämäki, Jyväskylä | 62.22814 | 25.64794 | Central Finland | 2018 | Male | W | x | 92938532 | 88633493 | 95.37 | 0.24 | 67242769 | 13.8414 | 92892355 | 98.88 | 0.25 | 91856037 | 14.0357 |
| **CAM015142** | Voiaskintie, Ulrikasund | 60.43466 | 25.32835 | Southern Finland | 2018 | Male | Y | x | 84316506 | 80894323 | 95.94 | 0.13 | 70140012 | 14.8534 | 84287184 | 99.04 | 0.14 | 83480957 | 15.0015 |
| **CAM015143** | Tvärminne, Hanko | 59.846 | 23.185 | Southern Finland | 2018 | Male | Y | x | 81341932 | 78370587 | 96.35 | 0.14 | 67261188 | 14.2685 | 81283183 | 99 | 0.15 | 80472992 | 14.4447 |
| **CAM015144** | Tvärminne, Hanko | 59.846 | 23.185 | Southern Finland | 2018 | Male | Y | x | 88405065 | 85421712 | 96.63 | 0.14 | 73193505 | 15.1229 | 88352550 | 98.97 | 0.15 | 87440926 | 15.3091 |

| | | | | | | | | | | | | | | | | | | | |
|---|---|---|---|---|---|---|---|---|---|---|---|---|---|---|---|---|---|---|---|
| **CAM015145** | Tvärminne, Hanko | 59.846 | 23.185 | Southern Finland | 2018 | Male | Y | x | 80894075 | 77989309 | 96.41 | 0.13 | 67679132 | 14.0029 | 80841879 | 98.97 | 0.14 | 80008747 | 14.195 |
| **CAM015146** | Tvärminne, Hanko | 59.846 | 23.185 | Southern Finland | 2018 | Male | Y | x | 81136931 | 78241764 | 96.43 | 0.13 | 67854265 | 14.2953 | 81086172 | 98.89 | 0.14 | 80187814 | 14.451 |
| **CAM015147** | Voiaskintie, Ulrikasund | 60.43466 | 25.32835 | Southern Finland | 2018 | Male | W | x | 75094177 | 72333859 | 96.32 | 0.13 | 62642058 | 13.4071 | 75042444 | 99 | 0.14 | 74295482 | 13.5672 |
| **CAM015148** | Voiaskintie, Ulrikasund | 60.43466 | 25.32835 | Southern Finland | 2018 | Male | W | x | 89834991 | 86506811 | 96.30 | 0.14 | 74333381 | 15.7749 | 89777636 | 99 | 0.15 | 88882799 | 15.9597 |
| **CAM015149** | Mosabackantie, Sipoo | 60.36973 | 24.54084 | Southern Finland | 2018 | Male | W | x | 73648890 | 71065467 | 96.49 | 0.13 | 61618406 | 13.0722 | 73604268 | 99.03 | 0.14 | 72886794 | 13.2296 |
| **CAM015150** | Voiaskintie, Ulrikasund | 60.43466 | 25.32835 | Southern Finland | 2018 | Male | W | x | 72378681 | 69440673 | 95.94 | 0.13 | 60321260 | 12.834 | 72332606 | 98.98 | 0.14 | 71594111 | 13.0016 |
| **CAM015151** | Voiaskintie, Ulrikasund | 60.43466 | 25.32835 | Southern Finland | 2018 | Male | W | x | 80232467 | 76779318 | 95.70 | 0.14 | 66143339 | 13.6987 | 80159712 | 99.06 | 0.15 | 79405646 | 13.9165 |
| **CAM015154** | Kanaküla 2 | 58.26328 | 25.11927 | Estonia | 2018 | Male | W | x | 71789797 | 69217313 | 96.42 | 0.13 | 60293429 | 12.7039 | 71746939 | 98.93 | 0.13 | 70982660 | 12.8702 |
| **CAM015155** | Kanaküla 1.2 | 58.15794 | 25.8694 | Estonia | 2018 | Male | W | x | 90309170 | 87203546 | 96.56 | 0.14 | 75335592 | 15.3226 | 90247219 | 98.99 | 0.14 | 89333141 | 15.5947 |
| **CAM015158** | Kanaküla 4 | 58.274533 | 25.117817 | Estonia | 2018 | Male | W | x | 77232567 | 74330844 | 96.24 | 0.14 | 64205532 | 13.123 | 82512306 | 98.99 | 0.14 | 76392613 | 13.3823 |
| **CAM015159** | Kanaküla 2 | 58.26328 | 25.11927 | Estonia | 2018 | Male | W | x | 79351392 | 76638691 | 96.58 | 0.13 | 66913224 | 13.7015 | 77179045 | 98.98 | 0.13 | 78483602 | 13.8975 |
| **CAM015162** | Thieves Hill, Aultmore, Keith | 57.575222 | -3.048778 | Scotland | 2015 | Male | Y | x | 82938651 | 79794024 | 96.21 | 0.15 | 67813099 | 14.2972 | 79287964 | 98.99 | 0.16 | 82016446 | 14.5358 |
| **CAM015163** | Portknockie, Buckie | 57.702785 | -2.879435 | Scotland | 2015 | Male | Y | x | 83269393 | 80197317 | 96.31 | 0.14 | 68632488 | 14.4383 | 82886426 | 98.95 | 0.15 | 82338256 | 14.6507 |
| **CAM015165** | Portknockie, Buckie | 57.702785 | -2.879435 | Scotland | 2015 | Male | Y | x | 71071794 | 47490 | 0.07 | 0.14 | 40949 | 0.00243509 | removed due to low mapping % | | | | |
| CAM015170 | Findlater Castle, Portsoy | 57.69157 | -2.77265 | Scotland | 2015 | Male | Y | x | 59557227 | 57359570 | 96.31 | 0.13 | 50112976 | 10.4116 | 71072090 | 0.79 | 0.13 | 58867297 | 10.5789 |
| **CAM015202** | F1 offspring of wild parents | NA | NA | Scotland | 2015 | Male | Y | x | 59937755 | 58011585 | 96.79 | 0.13 | 50720420 | 10.7925 | 59903009 | 99 | 0.13 | 59301604 | 10.9623 |
| **CAM015203** | F1 offspring of wild parents | NA | NA | Scotland | 2015 | Male | Y | x | 55068668 | 53331495 | 96.85 | 0.12 | 46913748 | 9.92293 | 57251863 | 99.03 | 0.13 | 54495189 | 10.0522 |
| **CAM015204** | F1 offspring of wild parents | NA | NA | Scotland | 2015 | Male | Y | x | 64822930 | 62348914 | 96.18 | 0.13 | 54148683 | 11.475 | 64780636 | 98.08 | 0.14 | 63535554 | 11.6182 |
| **CAM015206** | F1 offspring of wild parents | NA | NA | Scotland | 2015 | Male | Y | x | 87767164 | 84630216 | 96.43 | 0.16 | 70781662 | 14.6903 | 87719259 | 98.9 | 0.17 | 86758116 | 14.9508 |
| **CAM015207** | F1 offspring of wild parents | NA | NA | Scotland | 2015 | Male | Y | x | 62230737 | 58239265 | 93.59 | 0.12 | 51177246 | 10.9483 | 62197559 | 96.13 | 0.13 | 59791385 | 11.1071 |
| **CAM015208** | F1 offspring of wild parents | NA | NA | Scotland | 2015 | Male | Y | x | 47567413 | 46042450 | 96.79 | 0.12 | 40522208 | 8.60095 | 64848486 | 98.96 | 0.12 | 64176574 | 11.8856 |
| **CAM015209** | F1 offspring of wild parents | NA | NA | Scotland | 2015 | Male | Y | x | 119873564 | 114836831 | 95.80 | 0.17 | 95610964 | 19.8179 | 119796501 | 98.93 | 0.18 | 118513971 | 20.175 |
| **CAM015211** | F1 offspring of wild parents | NA | NA | Scotland | 2015 | Male | Y | x | 47171055 | 45865834 | 97.23 | 0.11 | 40747916 | 8.70432 | 63799250 | 99.04 | 0.11 | 63185074 | 11.9608 |
| **CAM15072** | Tvärminne, Hanko | 59.846 | 23.185 | Southern Finland | | Male | Y | | These samples were added to later analyses using the WW genome | | | | | | | | 0.08 | 71280369 | 14.1515 |
| **CAM015073** | Tvärminne, Hanko | 59.846 | 23.185 | Southern Finland | | Male | Y | | | | | | | | 68638162 | 99.02 | 0.08 | 67967326 | 13.4971 |
| **CAM015078** | Tvärminne, Hanko | 59.846 | 23.185 | Southern Finland | | Male | W | | | | | | | | 70063666 | 99.04 | 0.08 | 69389996 | 13.7484 |
| **CAM15084** | Tvärminne, Hanko | 59.846 | 23.185 | Southern Finland | | Male | W | | | | | | | | | | 0.08 | 70835562 | 13.9317 |

| | | | | | | | | | | | | | | | | | | | |
|---|---|---|---|---|---|---|---|---|---|---|---|---|---|---|---|---|---|---|---|
| **CAM015085** | Tvärminne, Hanko | 59.846 | 23.185 | Southern Finland | | Male | W | | | | | | | | 49250064 | 99.05 | 0.09 | 48782086 | 9.66459 |
| **CAM015086** | Tvärminne, Hanko | 59.846 | 23.185 | Southern Finland | | Male | Y | | | | | | | | 57402371 | 98.98 | 0.08 | 56815328 | 11.2504 |
| **CAM015088** | Tvärminne, Hanko | 59.846 | 23.185 | Southern Finland | | Male | Y | | | | | | | | 65556920 | 99.02 | 0.09 | 64913551 | 12.7004 |
| **CAM015089** | Tvärminne, Hanko | 59.846 | 23.185 | Southern Finland | | Male | W | | | | | | | | 67913274 | 99.02 | 0.09 | 67245909 | 13.2502 |
| **CAM015091** | Tvärminne, Hanko | 59.846 | 23.185 | Southern Finland | | Male | W | | | | | | | | 66279795 | 99.04 | 0.09 | 65642031 | 12.9068 |
| **CAM015094** | Tvärminne, Hanko | 59.846 | 23.185 | Southern Finland | | Male | Y | | | | | | | | 71591177 | 98.88 | 0.10 | 70791957 | 13.6097 |

Table S7: CRISPR sgRNAs sequences.

| **sgRNA name** | **Target sequence** | **Starting location in *valkea*** | **PAM** | **Specificity score %** | **Activity score** |
|---|---|---|---|---|---|
| Val1A | CTGAGAACACGGTACCTACG | WW_tarseq_419_arrow:7,014,923 | GGG | 62.5 | 0.68 |
| Val2A | ACACCCACAGTCTATTGCAG | WW_tarseq_419_arrow:7,021,659 | TGG | 62.5 | 0.505 |
| Val2B | TCGTGGCATTCTGCCGGCCG | WW_tarseq_419_arrow:7,021,588 | CGG | 62.5 | 0.698 |
| Val3A | CTCGGGGGAAGTATACACTT | WW_tarseq_419_arrow:7,031,836 | CGG | 83.3 | 0.722 |
| Val3B | CTTCGCTTACATCAGTGACA | WW_tarseq_419_arrow:7,031,950 | CGG | 83.3 | 0.754 |

Table S8: Elution gradient used in pheomelanin HPLC analysis.

| Time (s) | %A | %B |
|---|---|---|
| 0 | 96 | 4 |
| 0.2 | 96 | 4 |
| 0.3 | 94 | 6 |
| 20 | 94 | 6 |
| 21 | 60 | 40 |
| 35 | 40 | 60 |
| 39 | 40 | 60 |
| 42 | 96 | 4 |

Table S9: Waveform of disposable working electrode in pheomelanin analysis.

| Time (s) | Potential (V) | Integration |
|---|---|---|
| 0 | 0.13 | |
| 0.04 | 0.13 | |
| 0.05 | 0.45 | |
| 0.21 | 0.45 | Begin |
| 0.56 | 0.45 | End |
| 0.57 | -1.67 | |
| 0.58 | -1.67 | |
| 0.59 | 0.93 | |
| 0.6 | 0.13 | |
